# The somato-cognitive action network links diverse neuromodulatory targets for Parkinson’s disease

**DOI:** 10.1101/2023.12.12.571023

**Authors:** Jianxun Ren, Wei Zhang, Louisa Dahmani, Qingyu Hu, Changqing Jiang, Yan Bai, Gong-Jun Ji, Ying Zhou, Ping Zhang, Weiwei Wang, Kai Wang, Meiyun Wang, Luming Li, Danhong Wang, Hesheng Liu

## Abstract

The newly-recognized somato-cognitive action network (SCAN) is posited to be important in movement coordination. Functional disruptions in Parkinson’s disease (PD) correspond with complex, non-effector-specific motor symptoms, indicating that SCAN dysfunction may underlie these symptoms. Along the same lines, the SCAN may link multiple neuromodulatory targets used for PD treatment. To investigate the role of the SCAN in PD, we leveraged resting-state precision functional mapping, analyzing data from 673 individuals across 6 independent datasets and 5 types of neuromodulation. Our findings revealed functional abnormalities within the SCAN in PD patients and the selective involvement of the SCAN in diverse neuromodulatory targets. Moreover, our data suggests causal links between SCAN connectivity changes and motor symptom alleviation following both invasive and non-invasive neuromodulation. Collectively, these findings underscore the critical role of the SCAN in the pathophysiology of PD and its brain stimulation treatments, and suggest the SCAN as a promising candidate target for neuromodulation.

## Introduction

Several types of brain stimulation techniques are currently being employed as therapeutic interventions for Parkinson’s disease (PD). These include invasive approaches like deep brain stimulation (DBS) ^1^, as well as non-invasive techniques such as high-intensity focused ultrasound stimulation (FUS) ^2^, transcranial magnetic stimulation (TMS) ^3^, and transcranial electrical stimulation (tES) ^4^. These stimulations techniques target a diverse set of subcortical and cortical brain regions with the goal of alleviating PD symptoms, including the subthalamic nucleus (STN), globus pallidus pars internus (GPi), ventral intermediate nucleus of the thalamus (VIM) ^5^, supplementary motor area (SMA), and primary motor cortex (M1). The effectiveness of these treatment strategies in improving different aspects of motor symptoms varies, and the precise functional relationships among these neuromodulatory targets remain to be fully elucidated.

An impactful hypothesis proposed that both invasive and non-invasive stimulation targets for PD treatment may be linked by functional connectivity ^6^. This hypothesis has gained support stemming from the evidence that GPi- and STN-DBS targets exhibit highly similar functional networks ^7^, and that stimulation of the two targets induces comparable functional changes ^8^. Moreover, recent studies on brain stimulation mechanisms for PD highlighted that the stimulation of a cortical area or subcortical nucleus not only modulates its immediate vicinity, but also extends its influence to other brain regions within the same functional networks ^3,8–11^.

Recently, a new functional network, dubbed the somato-cognitive action network (SCAN), was unveiled. This network is thought to play a pivotal role in the integration of movement, and thus was suggested to be potentially relevant to PD ^12^. The SCAN is composed of a set of regions that interleaves effector-specific functional regions in the motor cortex, i.e. the canonical foot, hand, and mouth networks ^12^. These inter-effector regions differ from the effector regions in that they do not display a specificity for any body part. They were shown to be important in the integration of movement—planning and coordinating movement and executing axial body movements ^12^. Of particular interest, PD is associated with complex motor symptoms that correspond with disruptions in these functions ^13^, thereby pointing to SCAN dysfunction as a potential mechanism through which these symptoms are expressed.

The aim of the current study was to shed light on the role of the SCAN in PD. We leveraged the recently-established precision functional mapping approach—using large amounts of resting-state functional MRI (rsfMRI) data per individual—to answer three fundamental questions: (1) whether functional abnormalities exist within the SCAN in PD patients; (2) whether and how the SCAN may link existing neuromodulatory targets of PD; and (3) whether the SCAN plays a causal role in the modulatory effects of brain stimulation on clinical improvement. To probe these questions, we used three PD datasets comprised of extensively-sampled participants: a cross-sectional PD dataset (n = 166 PD patients, n = 65 healthy controls (HC), 30-minute rsfMRI scan for each participant), a STN-DBS dataset (n = 14 PD patients, incorporating 9 distinct combinations of stimulation conditions and follow-up timepoints, yielding a total of 300 minutes of rsfMRI scanning per participant; n = 25 HC, each with 19 minutes of rs-fMRI data), and a SMA-TMS dataset (n = 19 active TMS, n = 19 sham TMS, 17.4-minute rsfMRI scans for each participant). Moreover, we identified diverse neuromodulatory targets using a DBS sweet spot dataset in which 342 patients underwent STN-, GPi-, or VIM-DBS; a VIM-MRI guided FUS dataset with 10 patients with tremor-dominant PD, and a STN subregion segmentation dataset with 13 healthy participants.

## Results

### Functional abnormalities within the SCAN in PD patients

First, we sought to determine whether we could detect the SCAN in our cohorts. The SCAN was discernible in both healthy older individuals (Figure 1a and Extended Data Fig. 1a) and individuals diagnosed with PD (Figure 1b and Extended Data Fig. 1b). The SCAN motif, characterized by three distinct regions in the motor cortex, closely mirrors the spatial configuration identified by Gordon and colleagues ^12^. The cortical motif is also evident at the group level across healthy participants (n = 60, Figure 1c) and across PD patients (n = 65, Figure 1d). The PD patient group was subsampled from the full PD dataset (N = 166) to ensure equivalent demographics and imaging data quality relative to the healthy cohort (Extended Data Table 1).

**Fig 1.**
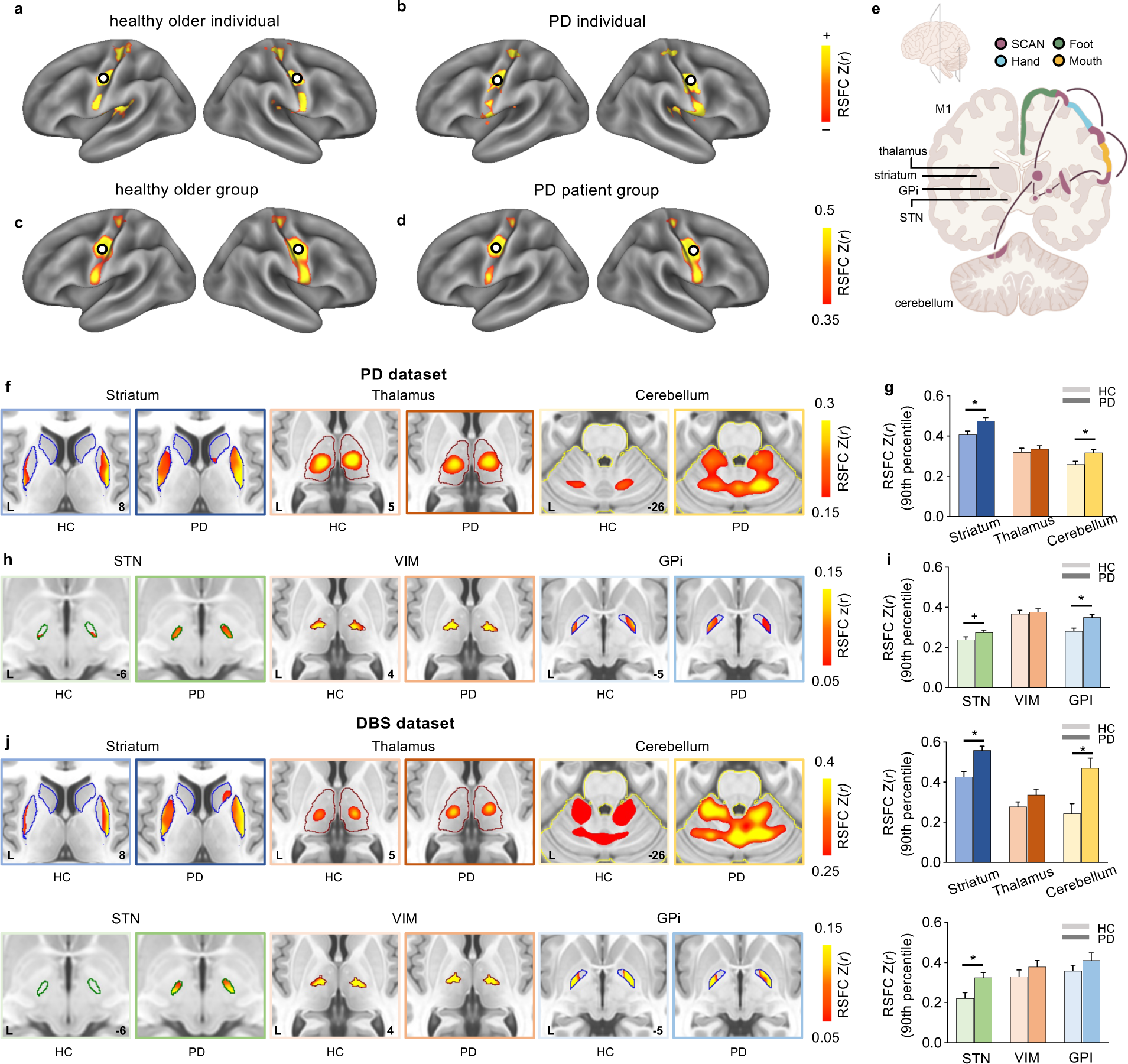
Abnormal hyper-connectivity between the SCAN and subcortical regions in PD patients. (a-d) The characteristic SCAN motif, consisting of three distinct regions within the M1 stripe, is observed consistently at the individual level in (a) healthy older individuals and (b) individual PD patients (randomly selected individuals are shown here), as well as at the group level in (c) a group of healthy older participants (n = 60) and (d) a group of PD patients (n = 65). (e) An illustration featuring coronal slices of the cerebrum and cerebellum shows the connectivity patterns between the SCAN (shown here interleaved with the foot, hand, and mouth effector regions) and subcortical regions, including the striatum, VIM, GPi, STN, and cerebellum. (f) In the PD dataset, the SCAN exhibits robust RSFC with the striatum (left), thalamus (middle), and cerebellum (right) in both healthy controls (HC) and PD patients, as depicted in axial views of these structures with color-coded structural boundaries. “L” denotes the left hemisphere and the numbers denote the axial slice indices. (g) The bar graph displays significantly higher RSFC between the SCAN and the striatum and cerebellum in PD patients relative to HC (*p < 0.05, FDR-corrected). (h) The SCAN also exhibits substantial RSFC with three nuclei targeted by DBS in PD, namely the STN (left), VIM (middle), and GPi (right). (i) Notably, PD displayed significantly greater or trends towards greater connectivity between the SCAN and the GPi (*p = 0.01, FDR- corrected) or STN (^+^p = 0.08, FDR-corrected). (j) These findings are corroborated in an independent dataset, the DBS dataset with 14 PD patients.

We then investigated the resting-state functional connectivity (RSFC) between the SCAN and subcortical regions that are involved in movement disorders ^12^. The SCAN exhibited robust RSFC with the striatum (putamen), thalamus, and cerebellum (Lobule VIII vermis) (striatum: RSFC = 0.41 ± 0.14, t(59) = 22.10, p < 0.001; thalamus: RSFC = 0.32 ± 0.15, t(59) = 16.48, p <0.001; cerebellum: RSFC = 0.26 ± 0.11; t(59) =17.46, p < 0.001; two-tailed one-sample t-tests, FDR-corrected). To determine the relationship between the SCAN and the neuromodulatory targets commonly used for PD treatment, we calculated the RSFC between the SCAN and the STN, GPi, and VIM. The SCAN showed significant RSFC with each neuromodulatory target (STN: RSFC = 0.24 ± 0.11, t(59) = 16.87, p < 0.001; VIM: RSFC = 0.37 ± 0.14, t(59) = 20.67, p <0.001; GPi: RSFC = 0.28 ± 0.12, t(59) = 18.04, p <0.001). All RSFC strengths and statistical results are summarized in Supplementary Table 1. A schematic illustration of the cortico-subcortical circuit associated with the SCAN is shown in Fig. 1e.

Compared to healthy participants, patients with PD displayed significantly greater or trends towards greater connectivity between the SCAN and the striatum, cerebellum, STN, and GPi (striatum: t(123) = 2.84, p = 0.01; cerebellum: t(123) = 1.98, p = 0.01; STN: t(123) = 1.93, p = 0.08; Gpi: t(123) = 3.36, p = 0.006; two-tailed two-sample t-tests, FDR-corrected) (Figure 1f-i), indicating abnormal hyper-connectivity in these brain regions in PD. However, no statistically significant differences were observed for the SCAN’s connectivity with the thalamus (t(123) = 0.66, p = 0.61, FDR-corrected) or the VIM (t(123) = 0.46, p = 0.65, FDR-corrected) (Figure 1f-i).

These findings were reaffirmed through replication analyses performed on the full PD dataset (n = 166), which yielded consistent conclusions (Extended Data Fig. 2). To test the generalizability of these findings, we extended our analyses to preoperative PD patients and healthy controls from the DBS dataset. Similar RSFC patterns and between-group statistical differences emerged in these additional analyses, apart from the lack of a significant difference between groups in SCAN-GPi RSFC (Figure 1j & Supplementary Table 1). The consistent observations coming from two independent datasets underscore the presence of functional abnormalities in SCAN connectivity among PD patients.

### Neuromodulatory targets for PD treatment selectively involve the SCAN

Recognizing the significant role of the SCAN in PD and its substantial connectivity with neuromodulatory targets, we investigated whether the SCAN is more highly functionally connected to precise neuromodulatory targets than the effector-specific networks are. To this end, we identified target regions from six datasets: 1) a STN target identified through volumes of tissue activated (VTA) by STN-DBS from the DBS dataset study ^10^; 2) the STN-DBS sweet spot estimated from 275 patients ^14^; 3) the motor subdivision of the STN (the ideal target of STN-DBS) ^15^; 4) the GPi-DBS sweet spot estimated from 28 patients ^14^; 5) the VIM-DBS sweet spot from 39 patients ^14^; and 6) the lesion overlap of MR-guided focused ultrasound lesioning of the VIM ^2^ (VIM-MRgFUS; see Extended Data Table 2 for more details). We calculated the whole-brain RSFC maps of the SCAN on the one hand, and of the three effector regions on the other hand, using the DBS dataset, and averaged each map’s RSFC within each of the six identified targets. We found that the RSFC map of the SCAN exhibited greater spatial overlap with each of the targets in comparison to the three effector-specific networks (Fig. 2). The RSFC between the SCAN and these targets was significantly greater than the effector regions’ in all cases (two-tailed paired-sample t-tests, all p’s < 0.01, FDR-corrected), signifying that PD neuromodulatory targets selectively involve the SCAN. This finding was replicated in the independent PD dataset, yielding concordant results (all p’s < 0.01, FDR-corrected; Extended Data Fig. 3), demonstrating the robustness of this finding. Additionally, to mitigate the potential confounding effect arising from the composite nature of the SCAN, which includes three distinct regions, in contrast to the single region of each effector-specific network, we compared the target RSFC with the SCAN against the target RSFC with the averaged three effector-specific networks. In both PD and DBS datasets, SCAN-target RSFC was significantly greater (all p’s < 0.01, FDR-corrected; Extended Data Fig. 4).

**Fig 2.**
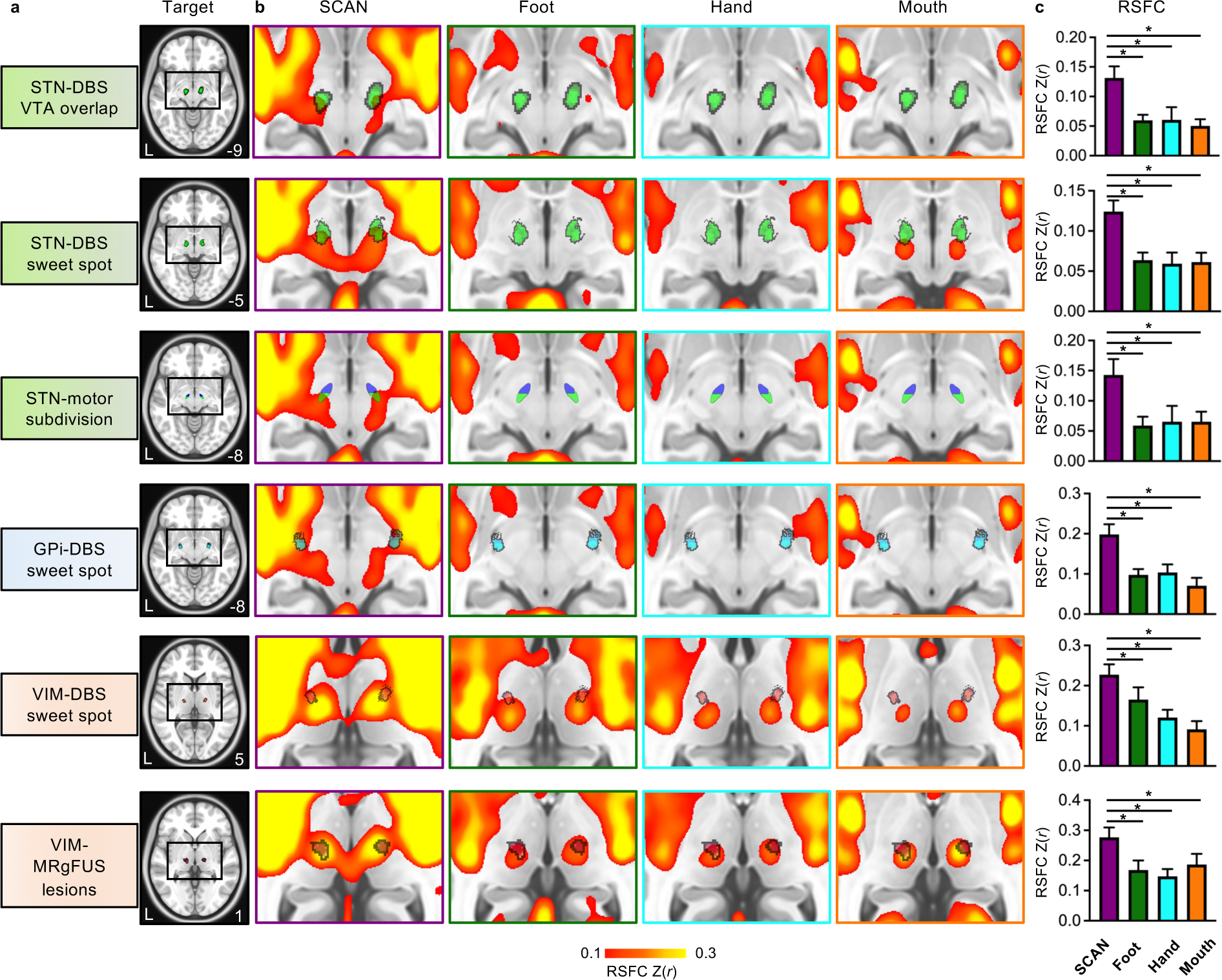
The SCAN displays greater functional connectivity with diverse neuromodulatory targets for PD treatment than primary motor effector regions. (a) Six distinct treatment targets for PD were identified from six datasets, including three STN targets (shown in green) identified from the STN-DBS VTA overlap, the STN-DBS sweet spot ^14^, and the motor subdivision of the STN^15^, a GPi target (shown in blue) from the GPi-DBS sweet spot ^14^, and two VIM targets (shown in red) from the VIM-DBS sweet spot ^14^ and the lesion overlap of VIM-MRgFUS from ^2^. An axial view is shown for each target using the 0.5-mm MNI ICBM 152 template. Black boxes delineate the areas used for the close-up views shown on the right, with the axial coordinate denoted on the bottom right. (b) Close-up views display group-averaged RSFC maps derived from the seeds in the SCAN (purple boxes), foot network (green boxes), hand network (blue boxes), and mouth network (orange boxes) across PD patients in the DBS dataset. The targets are overlaid on these RSFC maps. The SCAN RSFC maps exhibit greater overlap with the targets compared to the foot, hand, and mouth effector networks. (c) Bar graphs represent the average RSFC between each functional network and treatment target across PD patients in the DBS dataset. Each target is more highly connected to the SCAN (purple bars) than to the foot (green bars), hand (blue bars), or mouth (orange bars) functional networks (all p values < 0.01, FDR-corrected), indicating that they selectively involve the SCAN.

To further validate this finding, we performed an inverse RSFC analysis in the PD dataset, in which we calculated the group-averaged cortical RSFC maps of seed regions derived from the STN, GPi, and VIM sweet spots. Concordant with our results, the targets displayed cortical RSFC patterns all visibly involving the SCAN. Cumulatively, the average cortical RSFC map across the three targets predominantly coincided with the SCAN (Fig. 3a). Intriguingly, the target-based and average maps showed strong RSFC with the multiple regions of the cingulo-opercular network (CON), including the SMA, dorsal anterior cingulate cortex, and insula. These findings were replicated in the DBS dataset (Fig. 3b).

**Fig 3.**
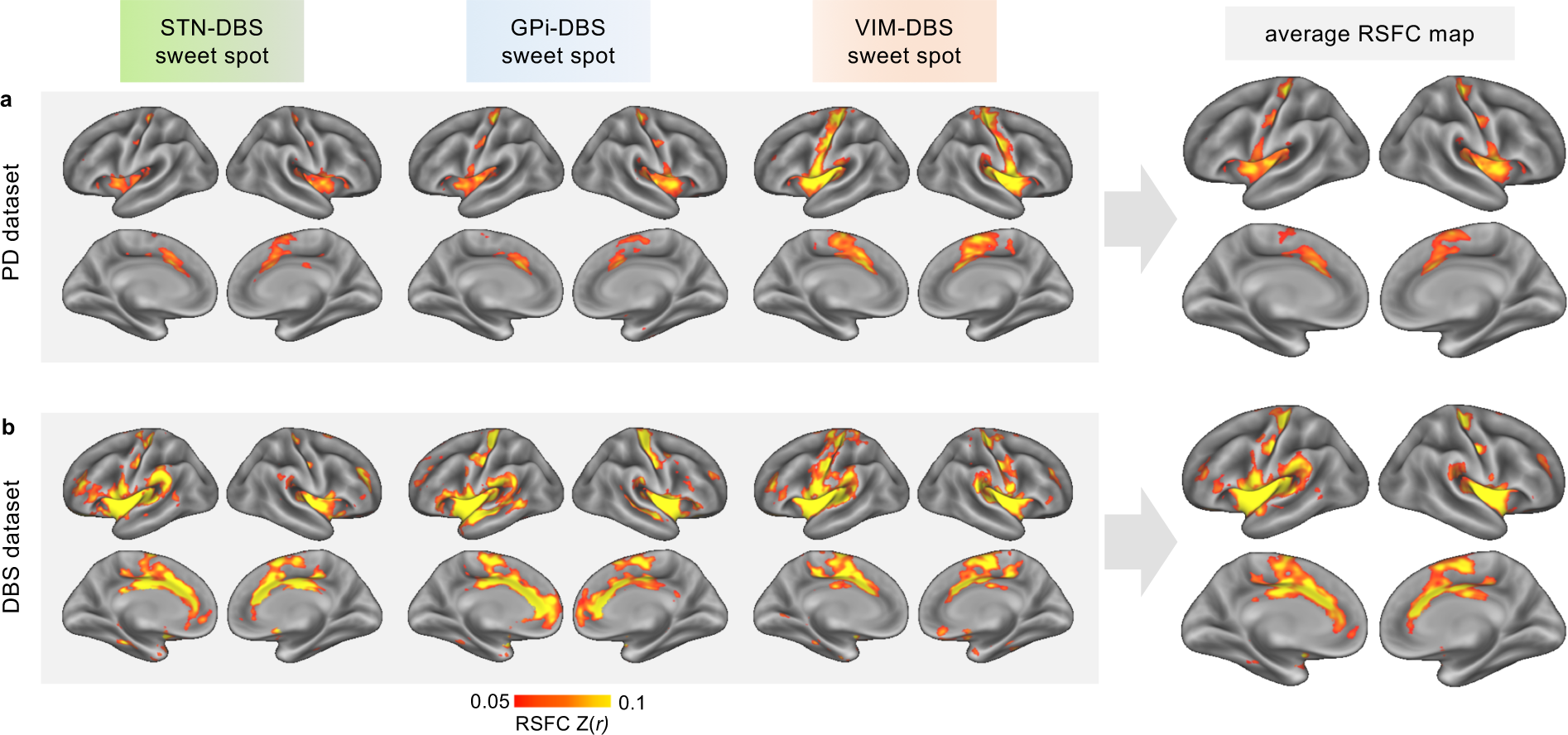
DBS sweet spots are functionally connected with the SCAN. (a) Group-averaged cortical RSFC maps were estimated based on the sweet spots of three DBS targets (STN, GPi, and VIM) across patients in the PD dataset. These three cortical RSFC maps exhibit connectivity to regions in M1 whose organization resembles the SCAN. Furthermore, these RSFC maps demonstrate strong connectivity with the cingulo-opercular network (CON), including the cingulate cortex and insula. The average RSFC map (the rightmost column) across RSFC maps derived from the three targets exhibits a clear pattern of connectivity with both the SCAN and the CON. (b) The RSFC maps were replicated in the DBS dataset.

In sum, this collection of evidence underscores the selective connectivity of routine PD neuromodulatory targets with the SCAN, hinting at the potential of stimulating the SCAN to relieve motor symptoms.

### RSFC changes in the SCAN are causally linked to motor symptom alleviation following brain stimulation

We examined whether brain stimulation of PD neuromodulatory targets modulates the SCAN, and whether its modulation is associated with motor symptom improvement. In the DBS dataset, 14 PD patients underwent bilateral STN-DBS electrodes implantation surgery (Fig. 4a; see detailed stimulation parameters in Supplementary Table 2). According to these parameters and electrode locations, we estimated the VTA for each patient; the VTA overlap mainly covered the STN (Fig. 4b). Motor symptom and precision rsfMRI data were collected before the surgery as well as for each stimulation state, i.e., ON or OFF. There were subsequent follow-up visits at 1, 3, 6, and 12 months. After receiving STN-DBS, significant improvements in motor symptoms were found in all participants (Fig. 4c, two-tailed paired sample t-tests, all p’s < 0.001, FDR-corrected). We investigated DBS-related changes in RSFC between the SCAN and the stimulation targets. With regards to STN VTA-SCAN RSFC, the preoperative RSFC in PD patients was notably higher than that observed in healthy controls (two-tailed two-sample t-test, t(37) = 2.55, *p* = 0.015). Across follow-up timepoints after implantation, reductions in RSFC emerged in comparison to the preoperative baseline, reflecting a “normalization” effect on the RSFC after DBS due to the downregulation of the STN VTA-SCAN hyperconnectivity. Furthermore, the DBS-ON RSFC was found to be mildly weaker than DBS-OFF RSFC in all follow-up periods, although these differences were not statistically significant (Fig. 4d, all p’s > 0.05, FDR-corrected). Additionally, the diminished VTA-SCAN RSFC in the ON vs. OFF state was found to correlate with improvement in total UPDRS-III scores, after adjusting for sex and age (Fig. 4e, partial Pearson correlation, *r* = 0.39, *p* = 0.007). RSFC change exhibited an association with various motor symptoms, including axial movement (Extended Data Fig. 5a, r = 0.44, p = 0.010, FDR-corrected), gait (Extended Data Fig. 5b, r = 0.39, p = 0.013, FDR-corrected), and bradykinesia (Extended Data Fig. 5c, r = 0.35, p = 0.023, FDR-corrected), but showed negligible correlations with tremor (Extended Data Fig. 5d, r = 0.04, p = 0.777, FDR-corrected) and rigidity (Extended Data Fig. 5e, r = 0.15, p = 0.303, FDR-corrected). These findings indicate that the SCAN-STN RSFC is causally linked to motor symptom alleviation during STN-DBS intervention, and that the DBS-related decrease in RSFC reflects a concurrent normalization of RSFC and the alleviation of motor symptoms.

**Fig 4.**
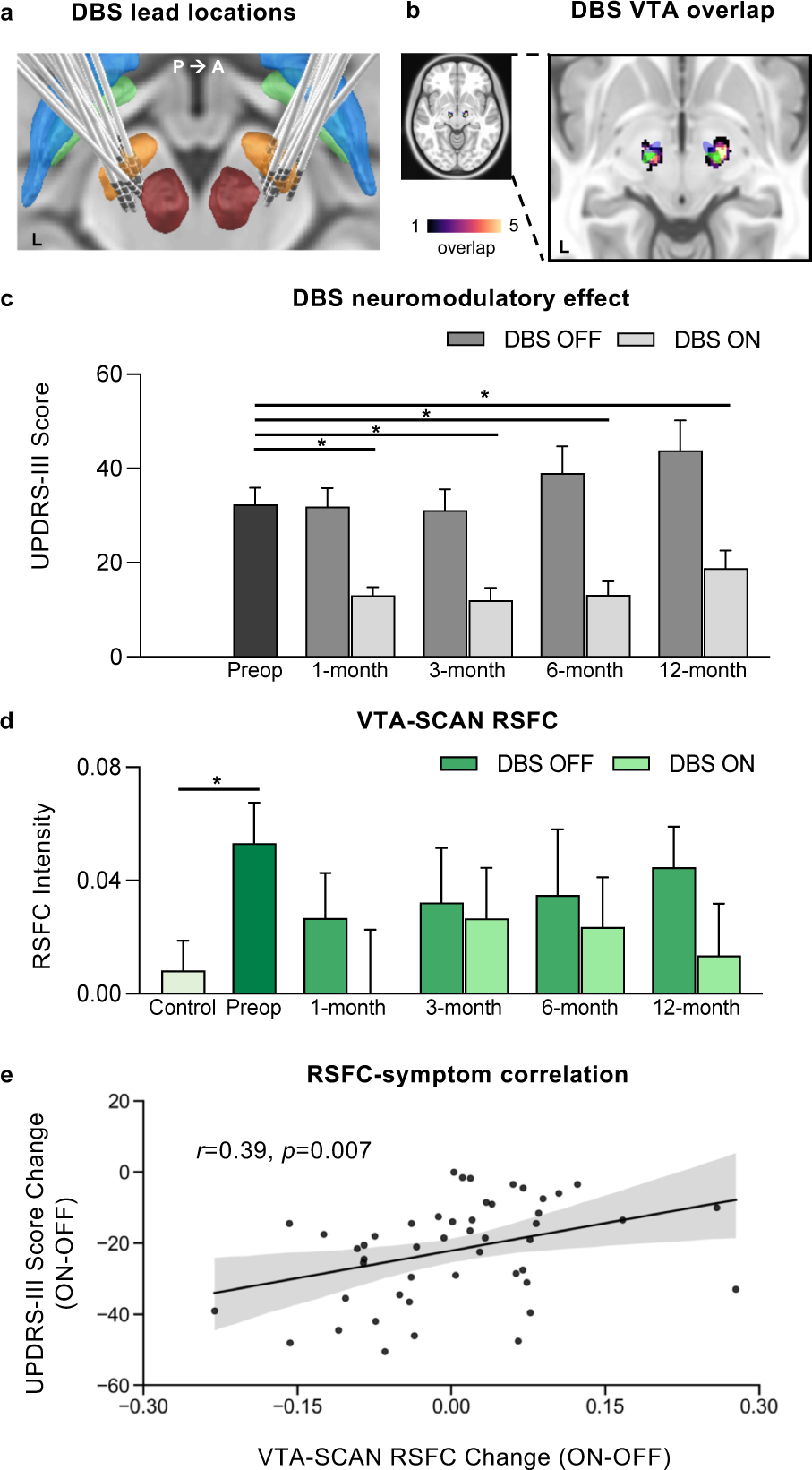
Changes in target-SCAN RSFC following STN-DBS are associated with motor symptom alleviation. (a) Electrode lead placements for all 14 PD patients in the DBS dataset and surrounding subcortical nuclei are presented. Bilateral STN (orange), GPe (blue), GPi (green), and the red nucleus (dark red) are shown. (b) The Volume of Tissue Activated (VTA) of STN-DBS was estimated for each patient and overlapped across all patients. The aggregated VTAs overlap primarily with the sensorimotor region of the STN (green). (c) The bar graph illustrates clinical outcomes assessed via UPDRS-III scores both preoperatively (preop, the black bar) and at 1, 3, 6, and 12 months post-STN-DBS surgery, in the DBS-OFF (dark gray bars) and DBS-ON (gray bars) states. The DBS-ON UPDRS-III scores are significantly lower compared to baseline assessments (all p’s < 0.01, FDR-corrected). (d) Preoperative VTA-SCAN RSFC among PD patients (dark green bar) is higher than that observed in healthy controls (light green bar, p = 0.013). Postoperative VTA-SCAN RSFC strength decreases relative to baseline. Moreover, the average RSFC is lower during DBS-ON than DBS-OFF, signifying a trend toward normalization as levels approach those observed in healthy participants. Error bars represent standard deviations. (e) The scatter plot shows the relationship between changes in VTA-SCAN RSFC and changes in symptoms experienced during transitions between the DBS-ON and DBS-OFF states, after adjusting for demographic variables (partial correlation, r = 0.39, p = 0.013). The shaded area represents the 95% confidence intervals. * p < 0.05.

In the TMS dataset ^3^, 38 PD patients were randomly allocated into two groups: one receiving active continuous theta-burst stimulation (cTBS) on the left SMA (n = 19), and the other undergoing sham cTBS (n = 19), on 14 consecutive days (Fig. 5a). After the treatment regimen, significant alleviation of motor symptoms, as evaluated by the total UPDRS-III scores, was evident in the active TMS group (Fig. 5b; pre = 27.05 ± 9.88, post = 18.95 ± 7.77; two-tailed paired-sample t-test, t(18) = -6.40, *p* < 0.0001). In contrast, the sham group exhibited no changes in motor symptoms (Fig. 5c; pre = 29.68 ± 8.90, post = 30.42 ± 9.66; t(18) = 0.85, p = 0.407). Given the relatively strong RSFC between the SMA and SCAN ^12^, it is plausible that stimulating the SMA could modulate SMA-SCAN RSFC. We found that the SMA-SCAN RSFC in PD patients is significantly weaker than that in aged healthy participants (Supplementary Fig.1; t(123) =2.17 , p = 0.032 in the PD dataset; t(37) = 2.37, p =0.023 in the DBS dataset). We assessed changes in SMA-SCAN RSFC before and after treatment, revealing a significant increase in RSFC strength within the active group (Fig. 5b; pre = 0.17 ± 0.09, post = 0.22 ± 0.11; t(18) = 2.23, p = 0.039), while the sham group exhibited no significant change (Fig. 5c; pre = 0.17 ± 0.14, post = 0.20 ± 0.14; t(18) = 0.49, p = 0.630). The increase in RSFC indicates a normalization effect of the active rTMS treatment. Furthermore, a significant correlation was found between change in SMA-SCAN RSFC and motor improvement in the active group but not the sham group, with adjustments for demographic factors (Fig. 5d; partial Pearson correlation, active: r = 0.47, p = 0.045; sham: r = - 0.35, p = 0.152). Of note, the scanning duration for each patient in the dataset might be too short to draw definitive conclusions (see Limitations), but these results are consistent with those from the DBS dataset.

**Fig 5.**
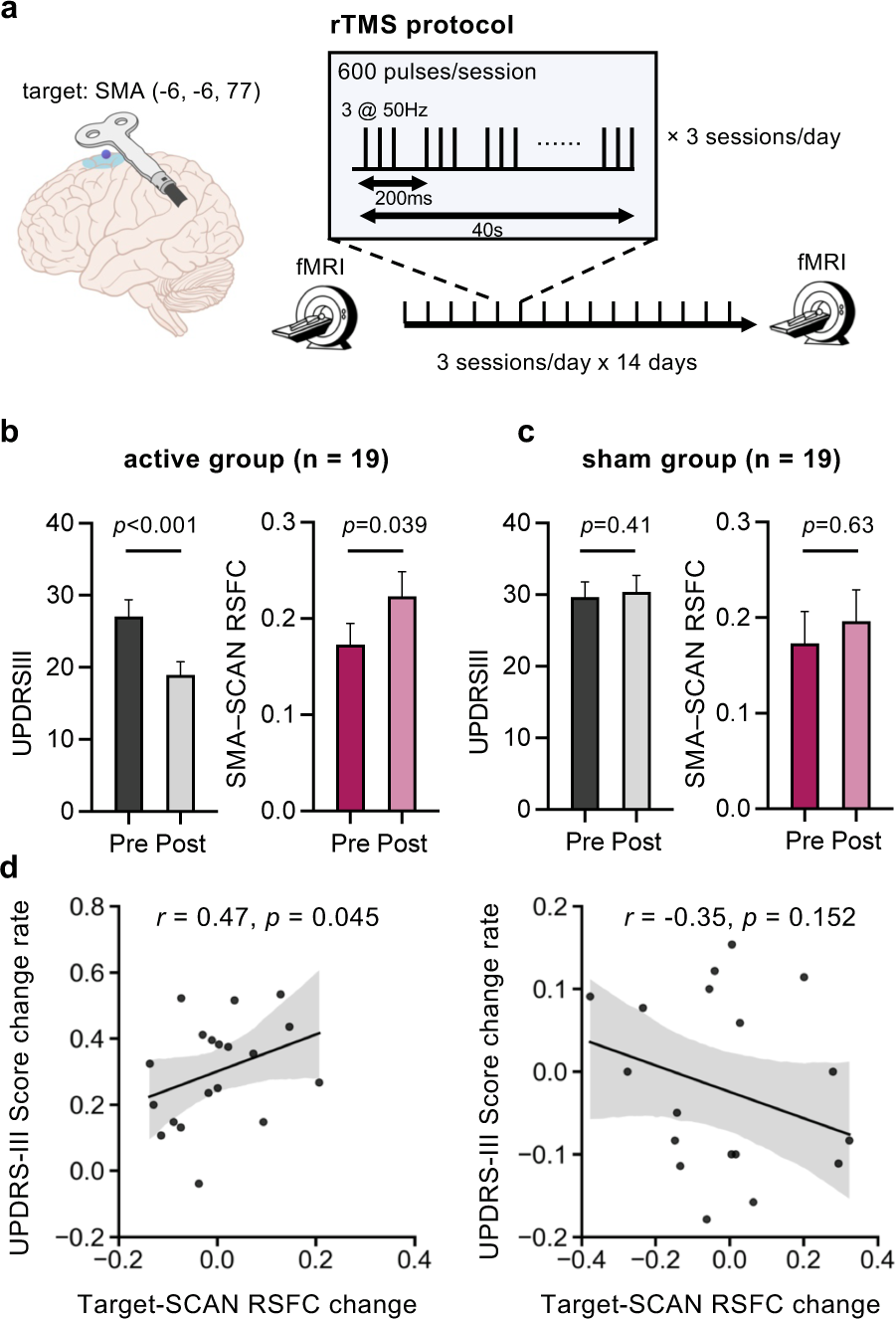
The normalization of target-SCAN RSFC following SMA-rTMS is associated with motor symptom alleviation. (a) The left illustration shows the left SMA target region for rTMS and its MNI coordinates (-6, 6, 77). PD327 patients underwent 14 consecutive days of either active or sham continuous Theta Burst Stimulation (cTBS), with three sessions of 600- pulse cTBS administered each day at 15-minute intervals. Rs-fMRI scans were conducted before and after the treatment. (b) In the active treatment group, a significant alleviation of motor symptoms as measured with the UPDRS-III was observed (p < 0.001), accompanied by a significant increase in target-SCAN RSFC (p = 0.039). This increase reflects a normalization effect, with stronger RSFC values approaching those observed in healthy participants (see Supplementary Fig. 1). (c) In the sham treatment group, both motor symptoms and target-SCAN RSFC remained relatively unchanged (p > 0.05). (d) The scatter plot in the active treatment group shows a significant correlation between changes in target-SCAN RSFC and changes in UPDRS-III scores, whereby larger increases in RSFC are associated with greater symptom improvement, adjusting for demographic variables (left panel, partial correlation, r = 0.47, p = 0.045). This association was not present in the sham group (right panel, r = -0.35, p = 0.152).

Collectively, these findings establish a causal link between target-SCAN RSFC and motor symptoms in PD, through both invasive and non-invasive brain stimulation approaches. This further highlights the critical role of the SCAN in the context of brain stimulation treatments for PD (Extended Data Fig. 6).

## Discussion

In this study, we used multiple extensively-sampled precision rsfMRI datasets to investigate the SCAN, a newly recognized functional network closely tied to movement coordination, in the context of PD. Our analyses yielded several key insights. First, we revealed reproducible functional abnormalities in RSFC between the SCAN and subcortical regions in PD patients. Second, the SCAN exhibited stronger functional connectivity with diverse PD subcortical neuromodulatory targets compared to the motor cortex’s effector regions. Third, our investigations across both invasive and non-invasive brain stimulation studies revealed causal links between changes in target-SCAN RSFC and motor symptom improvement. Overall, these findings underscore the significance of the SCAN in PD, and suggest that the SCAN may be selectively modulated in multiple PD brain stimulation treatments, including DBS, MRgFUS, and rTMS. Importantly, the SCAN emerges as a promising candidate target for neuromodulation.

Our study represents a timely clinical translation of novel neuroscience discoveries that enhance our comprehension of network-level pathophysiology and neuromodulatory mechanisms in PD. In our investigation, we unveiled functional abnormalities within a newly-recognized cortico-subcortical circuit implicated in PD. Additionally, our observations confirm that the commonly targeted regions for neuromodulation selectively involve this circuit. This finding is an extension of prior conclusions that highlight a highly similar brain network as being engaged following STN- and GPi-DBS stimulation^16^, supporting the hypothesis that various brain stimulation targets may share a common functional network^17^. Importantly, our results indicate that stimulating these targets alleviates PD symptoms by normalizing abnormal functional connectivity patterns, in line with prior studies ^2,9^.

Our results also suggest that the SCAN is a promising candidate target for neuromodulation. We demonstrated that the optimal stimulation sites for various DBS targets exhibit highly similar connectivity patterns, which are selectively linked to the SCAN. Moreover, we established a causal relationship between target-SCAN RSFC and clinical outcomes. These findings underscore an important role of the SCAN in current neuromodulatory therapies, and provide support for directly or indirectly targeting the SCAN to achieve improved clinical efficacy. On one hand, the SCAN may be employed to personalize the localization of DBS targets ^18,19^. For instance, GPi-DBS yields substantial variability in clinical outcomes, which may be attributed to individual differences in the functional organization of the GPi ^20^. Targeting the strongest individualized functional projection in the GPi from the SCAN using precision fMRI may improve targeting accuracy and enhance treatment response rates. On the other hand, the integration of functional neuroimaging and machine learning algorithms exhibits a promising avenue for automating DBS programming ^21^. The incorporation of the cortical-subcortical circuit of the SCAN in the procedure may further refine the accuracy of functional imaging-assisted programming.

The SCAN is localized on the surface of the cortex, and therefore is accessible to non-invasive brain stimulation, such as rTMS. Despite extensive explorations of rTMS for PD treatment, clinical outcomes remain suboptimal ^22^. Recent research in other brain disorders, such as depression and post-stroke aphasia, suggests that inaccurate target localization may contribute to these unsatisfactory results ^23–26^. Conventionally, rTMS targets for PD include the primary motor area, often described broadly as “M1” or more specifically as the hand/foot region within M1^27^. We found partial overlap between the electric field (E-field) induced by M1-rTMS and the SCAN (Supplementary Fig. 2a). This observation raises the question of whether it is the partial stimulation of the SCAN, rather than effector-specific regions, that is responsible for the observed clinical improvement. Another common rTMS target is the SMA, which also exhibits E-field overlap with the SCAN (Supplementary Fig. 2b). Beyond this overlap, the SMA, as a critical node in the CON, possesses strong connectivity with the SCAN and is thought to mediate motor planning, preparation, and execution ^12,28,29^. Our results also illustrated that SMA-rTMS normalized RSFC between the SMA and SCAN, indirectly modulating the SCAN. Additionally, studies employing direct cortical stimulation for alleviating parkinsonism show discordant results^30,31^, similar to rTMS, which may also partially be due to variations in stimulation location within the motor cortex. Woolsey and colleagues reported two intriguing cases of patients with severe PD or with parkinsonism, in whom motor symptoms strikingly disappeared following direct cortical stimulation of the motor cortex^31^. The symptom disappearance may be more straightforwardly explained as a consequence of SCAN stimulation, rather than stimulation of effector-specific regions. The alleviation of motor symptoms, however, has not been replicated through subdural motor cortical stimulation in a cohort of five patients with refractory parkinsonism^30^. Due to the limited areal size of each inter-effector region in the SCAN, delivering stimulation effectively to the network without precise localization may be challenging, potentially yielding variable responses. Consequently, future applications involving direct, personalized targeting of the SCAN through neuronavigated rTMS may offer a more effective means of modulating the specific cortico-subcortical circuit, potentially yielding enhanced clinical response in PD ^32^ (Extended Data Fig. 6). Directly targeting the SCAN may also be promising for other movement disorders characterized by disturbances in motor coordination, such as essential tremor and dystonia.

Our study indicates that the SCAN has a functional role distinct from that of other effector-specific networks. Despite substantial evidence that the SCAN lacks movement specificity in the original study ^12^, the functional role of the SCAN remains a subject of debate ^33^. An alternative viewpoint posits that it may mediate specific movement functions associated with the abdomen, upper face, and throat^33^. PD serves as an appropriate disease model to address this debate since it predominantly manifests as disruptions in nonspecific motor functions, such as bradykinesia, tremor, and rigidity ^13^. Our study, having revealed SCAN functional abnormalities in PD, supports the former notion, however further validation is needed. It is also important to take into consideration the subcortical structures that are connected to the SCAN. The basal ganglia, including the GPi and STN, and thalamus, including the VIM, hold pivotal roles in information integration, including motor integration ^20,34–37^. Importantly, they exhibit stronger connectivity with the SCAN in comparison to effector-specific networks (Fig. 3). Collectively, these regions and networks may constitute an integrated cortico-basal-ganglia-thalamo-cerebellar circuit ^36^ (Fig 1e) that governs motor integration, coordination, and planning. This integrated action control system is less likely to be specialized for regulating specific motor actions, and more likely to have a role in the broader context of motor function regulation.

From a technical perspective, the current study demonstrates the importance of employing precision fMRI when investigating brain disorders. To uncover reproducible findings, it is imperative to lengthen the duration of fMRI acquisition to obtain reliable individualized functional imaging data. The typical 6-minute rs-fMRI scans routinely collected in the RSFC literature have relatively low reliability due to limited temporal signal-to-noise ratio, making it harder to detect brain-phenotype associations ^38^. Additionally, it is essential to adopt personalized fMRI techniques to capture precise functional organization information at the individual level ^39^. The conventional approach of employing group-based methods to mitigate the influence of imaging noise leads to the blurring of fine-grained functional features ^39–42^. Despite the growing use of precision fMRI in healthy populations and the important insights that have emerged from such studies ^40,43,44^, the acquisition of longer fMRI sessions in patients is regarded as challenging due to their lower compliance and increased motion inside the scanner. In our study, we were able to acquire over 30 minutes of rs-fMRI data in hundreds of PD patients, demonstrating the feasibility of performing precision fMRI in challenging patient populations. This enabled us to unveil reproducible functional abnormalities, selective involvement of the SCAN in neuromodulatory therapies, and causal effects of SCAN neuromodulation across independent PD datasets. These outcomes underscore the significance of employing precision fMRI as a tool for investigating brain disorders.

Several limitations warrant consideration in our study. First, the medication state differed between the PD and DBS datasets: patients in the PD dataset underwent scanning in the medication ON state, while patients in the DBS dataset did so in the medication OFF state. However, the fact that the findings were closely replicated in the two datasets demonstrates that the medication state did not drive the results. Second, the scanning duration in the TMS study was relatively short, with a duration of 17.4 minutes for each patient. Consequently, the outcomes derived from the TMS dataset should be regarded as preliminary, necessitating further validation in studies with more extensive rsfMRI data per patient. Finally, while our study implies that the SCAN may hold promise as a non-invasive brain stimulation target for PD treatment, whether direct targeting of the network can yield lasting improvements in motor symptoms remains to be determined. As such, efficacy and safety of directly targeting the SCAN are to be prospectively evaluated in randomized clinical trials. Moreover, considering the representation of three distinct inter-effector regions in the SCAN, it is worth assessing whether targeting each of them elicits equivalent or distinct response profiles.

In summary, the SCAN may play a pivotal role in PD, as it exhibits substantial functional abnormalities, is selectively involved in neuromodulatory therapies employing diverse targets, and demonstrates casual links with clinical improvement. The SCAN is a testable candidate neuromodulatory target for PD treatment and warrants evaluation in future clinical trials.

## Methods

In the study, we used six independent datasets with 673 participants, comprising 1) a PD dataset of 166 PD patients and 65 healthy controls, 2) a DBS dataset featuring 14 PD patients, with evaluations conducted both pre- and post-DBS surgery, along with 25 healthy controls, 3) a TMS dataset involving 38 PD patients, 4) the DBS sweet spot dataset with 342 patients, and 5) the STN subregion segmentation dataset with 13 participants, as well as 6) the VIM-MRgFUS dataset, which includes 10 patients with tremor-dominant PD. The following sections provide detailed descriptions of each dataset regarding their inclusion and exclusion criteria, participant demographics, and the acquisition of imaging data.

### Dataset 1: PD dataset

#### Patients

A total of 180 PD patients were recruited from Henan Provincial People’s Hospital, China. The inclusion criteria included being aged 18 years or above and a confirmed diagnosis of idiopathic PD. Exclusion criteria comprised the following: 1) MRI contraindications; 2) a history of neurological disorders aside from PD, including stroke, cerebrovascular disease, seizures, and brain tumors; 3) prior invasive neurosurgeries such as DBS or ablation; and 4) average relative head motion larger than 0.2 mm during rsfMRI scanning. Four patients did not complete MRI scanning, and 10 patients were excluded due to excessive head motion. Ultimately, 166 patients were included in the analysis (64 women, 102 men; mean age = 61.8 years, SD = 7.84; see demographic and clinical details in Table 1).

#### Healthy controls

71 healthy participants aged 18 years or older, devoid of neurological or psychiatric disorders, were enrolled. Exclusion criteria included MRI contraindications and an average relative head motion exceeding 0.2 mm. After excluding 11 participants due to excessive head motion, the analysis included 60 healthy control participants (34 women, 26 men; mean age = 56.10 years, SD = 6.64; see Table 1). The control group exhibited significantly different demographics from the PD group. We thus sampled a subset of 60 PD patients from the 166 patients to ensure demographic matching when performing case-control analyses (Extended Table 1). Ethics approval for the data collection was obtained from the Henan Provincial People’s Hospital Institutional Review Board. Written informed consent was obtained from all participants. ***Imaging acquisition.*** Participants underwent a series of MRI procedures, including one structural MRI scan lasting 8 minutes and 50 seconds, and five rs-fMRI scans, each spanning 6 minutes and 14 seconds, resulting in a cumulative scan duration of 31 minutes and 10 seconds. All MRI acquisitions were performed using a Siemens 3T Prisma MRI scanner equipped with a 64-channel head coil. The structural scans involved T1-weighted images acquired through a MP2RAGE sequence (TI1 = 755 ms, TI2 = 2500 ms, TE = 3.43 ms, TR = 5,000 ms, flip angle1 = 4°, flip angle2 = 5°, matrix size = 256 ×256, 208 sagittal slices, spatial resolution = 1 × 1 × 1 mm^3^). An acceleration factor of 3 (with 32 reference lines) was applied in the primary phase encoding direction, with online GRAPPA image reconstruction. Rs-fMRI data were acquired using an gradient-echo echo planar imaging (GE-EPI) sequence (TE = 35 ms, TR = 2,000 ms, flip angle = 80°, and 75 slices, spatial resolution = 2.2 × 2.2 × 2.2 mm^3^). During data acquisition, participants were instructed to maintain open eyes, remain awake while keeping their body still, and minimize head movement.

### Dataset 2: DBS dataset

#### Patients

A total of 14 patients (mean age: 54.71 ± 7.65 years, age range: 40 to 67 years, 5 women, 9 men) diagnosed with the akinetic-rigid dominant form of idiopathic PD were recruited from three centers, including Tiantan Hospital, Beijing; Peking Union Medical College Hospital, Beijing; and Qilu Hospital, Jinan, China. Inclusion criteria consisted of: 1) age between 18 and 75 years; 2) Mini-Mental State Examination (MMSE) score above 24; 3) Hoehn and Yahr scale (H-Y) above stage two in the medication OFF status; 4) PD duration exceeding five years; 5) established positive response to dopaminergic medication (at least 30% UPDRS-III improvement with levodopa); and 6) ability to provide informed consent, assessed through preoperative neuropsychological evaluation. Exclusion criteria encompassed: 1) ineligibility for DBS, such as anesthesia complications; 2) history of hydrocephalus, brain atrophy, cerebral infarction, or cerebrovascular diseases; 3) inability to comply with verbal instructions; 4) presence of severe chronic conditions that could confound data interpretation; 5) MRI contraindications or inability to complete MRI scans. Out of the initial cohort, 11 patients had a complete dataset, while three patients had incomplete data due to missing post-surgical visits (DBS01 after the 1-month follow-up, DBS03 after the 3-month follow-up, and DBS08 at the 1-month follow-up only). No adverse events were reported during the study. The clinical trial was registered on ClinicalTrials.gov under the identifier NCT02937727. Ethics approval for this project was granted by the ethics committees of Tiantan Hospital, Peking Union Medical College Hospital, and Qilu Hospital. Written informed consent was obtained from all participating individuals.

Each patient underwent standard frame-based stereotaxic DBS implantation surgery at one of the aforementioned medical institutions. The bilateral STN were the targeted regions for DBS, localized through presurgical structural MRI scans, intra-operative electrophysiological recordings, and observed motor symptom improvement during the surgery. Two quadripolar DBS electrodes (Model L301C, Pins Medical Co., Beijing, China) were bilaterally implanted into the STN for each patient. A low field potential sensing-enabled neurostimulator (G106R, Beijing Pins Medical Co., Ltd) was connected to the leads (Model E202C, Pins Medical Co., Beijing, China) during a single operation. The DBS stimulator and electrodes were compatible with the 3T MRI environment and proven safe for MRI scans with implantation. At each postsurgical visit, a team of two neurologists managed each patient’s DBS system. Optimized DBS programming, resulting in optimal motor symptom improvement, was achieved by selecting positive and negative contacts and determining stimulation frequency, amplitude, and pulse width.

#### Healthy controls

Additionally, healthy control participants matched in age to the patient group were recruited. Similar exclusion criteria were applied, encompassing relevant medical history, ability to follow instructions, conditions that could complicate data interpretation, MRI contraindications, and average relative head motion exceeding 0.2 mm. The control group comprised 28 participants. One participant was excluded due to incomplete MRI data caused by discomfort in the scanner, and two participants because of excessive head motion, leaving 25 participants suitable for the case-control analysis (Extended Table 1; 13 women and 12 men; mean age = 56.32 ± 6.88).

#### Imaging acquisition

Participants underwent data acquisition across five visits, including one presurgical and four post-surgical follow-up visits. The presurgical visit occurred approximately one month before the DBS surgery, while the postsurgical visits occurred at 1, 3, 6, and 12 months after surgery. Data acquisition encompassed MRI scans, neurological assessments, and CT scans. Of note, the presurgical visit involved T1w MRI run and 5 rs-fMRI runs (totaling 31 minutes of rsfMRI). For each postsurgical visit, participants underwent four runs of DBS ON (130-Hz continuous stimulation) fMRI (25 minutes) followed by four runs of DBS OFF fMRI (25 minutes). Control participants attended one visit, involving one T1-weighted MRI run and three BOLD fMRI runs lasting 19 minutes in total.

All MRI data were collected using a 3T Philips Achieva TX whole-body MRI scanner equipped with a 32-channel head coil. T1-weighted structural images were acquired using a MPRAGE sequence, lasting 4 minutes and 14 seconds (TE=3.70 ms, TR = 7.52 ms, flip angle=8°, 180 sagittal slices, spatial resolution = 1 × 1 × 1 mm³). Functional images were acquired with a 6- minute and 14-second transversal GE-EPI sequence (TE=30 ms, TR=2000 ms, flip angle=90°, 37 slices, spatial resolution = 2.875 × 2.875 × 4 mm³, 184 frames/run). Computerized tomography (CT) images were acquired with a uCT 760 (United Imaging, Shanghai) scanner one month after surgery. A head helical sequence, with FOV=512×512, pixel spacing=0.449 mm × 0.449 mm, 204 slices, slice thickness=0.625 mm, was used.

### Dataset 3: TMS dataset

#### Patients

The TMS dataset was previously documented in a randomized clinical trial paper ^3^. Enrolment of participants occurred at the First Affiliated Hospital of Anhui Medical University. Inclusion criteria encompassed: (a) confirmed diagnosis of idiopathic PD; (b) stable medication treatment for a minimum of 2 months; (c) age of 40 years or older; and (d) MMSE score surpassing 24. Exclusion criteria included: (a) history of addiction, psychiatric disorders, or neurological conditions apart from PD; (b) discernible focal brain lesions on T1-/T2-weighted fluid-attenuated inversion recovery images; (c) modifications in anti-PD medications during rTMS; (d) substance abuse within the preceding 6 months; (e) presence of nonremovable metal objects near or within the head; (f) previous experience with rTMS treatment; and (g) a history of seizures or familial history of seizures in first-degree relatives. Review and approval of the study protocol were carried out by the institutional ethics committee at the First Affiliated Hospital of Anhui Medical University. All participants granted informed consent in written form prior to engaging in the study. The study was prospectively registered on ClinicalTrials.gov under the identifier NCT02969941.

Among the cohort, 42 patients were randomly divided into two groups, with each group receiving either active (n = 22) or sham (n = 20) continuous theta-burst stimulation (cTBS) over a span of 14 consecutive days. On each treatment day, three sessions of 600-pulse cTBS were administered with 15-minute intervals. The stimulation was directed towards the left SMA proper with MNI coordinates of −6, −6, 77, guided by the neuronavigation system (Brainsight; Rogue Research, Montreal, QC, Canada), and applied at 80% of the resting motor threshold (RMT). The active group underwent TMS using a Magstim Rapid2 transcranial magnetic stimulator (Magstim Company, Whitland, UK) equipped with a 70-mm air-cooled figure-of-eight coil. Conversely, the sham group received treatment using a placebo coil (Magstim) designed to simulate the sensory experiences and auditory cues of the actual stimulation.

#### Imaging acquisition

Both structural and functional MRI data were obtained using a 3-T scanner (Discovery 750; GE Healthcare, Milwaukee, WI). High-resolution T1-weighted structural images were captured using a three-dimensional brain-volume sequence (TE = 3.18 ms, TR = 8.16 ms, flip angle = 12°, 188 sagittal slices, voxel size = 1 × 1 × 1 mm³). Functional images were obtained through a single-shot gradient-recalled EPI sequence (TE = 30 ms, TR = 2400 ms, flip angle = 90°, 46 transverse slices, voxel size = 3 × 3 × 3 mm³). A total of 217 functional image frames were collected from each participant, equivalent to approximately 8.68 minutes, both prior to and post TMS treatment. Prior to scanning commencement, participants were instructed to maintain a resting state with their eyes closed, ensuring wakefulness while keeping the body still and minimizing any head movement. In the current study, both the active and sham group each had one patient who did not complete the rs-fMRI scanning. Additionally, two patients from the active group and one patient from the sham group were excluded from the analysis due to average relative head motion greater than 0.2 mm, leaving 19 patients in each group (Extended Table 1).

### Neuromodulation datasets

Multiple neuromodulatory targets were consolidated from studies which identified regions of interest (ROIs) from a meta-analysis of DBS sweet spots^14^, VTA overlap derived from the DBS dataset^10^, segmentation of the STN^15^, and lesion overlap from a VIM-MRgFUS dataset^2^. We generated the target ROIs by applying a binary transformation to the probabilistic values of the sweet spots or overlap maps with thresholds greater than zero.

### DBS sweet spots dataset

Elias et al. ^14^ conducted a comprehensive retrospective multicohort study involving DBS. In this dataset, there were 275 patients who underwent STN-DBS (80 women, 195 men, mean age = 59.8 ± 7.1 years), 28 patients with GPi-DBS (13 women, 15 men, mean age = 64.4 ± 7.0 years), and 39 patients with VIM-DBS (13 women, 26 men, mean age = 64.3 ± 11.6 years). Using probabilistic stimulation mapping, they generated sweet spot atlases for STN-, GPi-, and VIM-DBS. The specific atlas employed was the pre-installed version within the LEAD-DBS software.

### VTA overlap of the DBS dataset

We generated the VTA overlap of the DBS dataset as an additional STN-DBS target atlas (see the DBS electrode localization and the VTA estimation subsection).

### STN segmentation dataset

Accolla et al. ^15^ performed the STN segmentation using diffusion-based tractography derived from data of 13 participants (6 women, 7 men, mean age = 50.6 ± 10.9 years, age range: 40-72 years). The STN underwent segmentation into three distinct subregions, including the sensorimotor, associative, and limbic subregions. The sensorimotor subregion is regarded as the ideal target for the STN-DBS. The atlas used was the pre-installed version in the LEAD-DBS software ^45^.

### VIM-MRgFUS dataset

In a prior investigation, Dahmani et al. ^2^ recruited a cohort of 10 patients with tremor-dominant PD who underwent VIM-MRgFUS treatment (2 women, 8 men, mean age = 55.4 ± 7.2 years). Substantial alleviation of tremor symptoms was observed in all patients. The MRgFUS lesions were manually delineated by a radiologist and subsequently overlapped to generate a lesion overlap map of the VIM target.

### UPDRS assessments

The primary outcome measure of patients’ motor symptoms was assessed using the Unified Parkinson’s Disease Rating Scale-III (UPDRS-III). In the PD dataset, the Movement Disorder Society (MDS)-sponsored revision of the UPDRS ^46^ was conducted to evaluate PD motor symptoms in the medication ON condition. In the DBS dataset, comprehensive evaluations were recorded using the original version of the UPDRS-III during a medication OFF state, with patients refraining from taking medication for a minimum of 12 hours. Subsequently, two experienced neurologists independently scored each UPDRS-III subitem based on the recorded video material. Rigidity-related subitems were assessed by an on-site neurologist. These assessments exhibited substantial inter-rater reliability (ICC = 0.90)^10^. The scores employed in this study represent the averages of the two assessors’ scores. To account for the differences between the two UPDRS versions, a simplified conversion approach was implemented to yield adjusted UPDRS-III total scores, by subtracting seven points from the MDS UPDRS-III total scores, following the methodology reported in a prior publication ^47^. In the TMS dataset, the original version of the UPDRS-III evaluations took place in the medication OFF state, both prior to and after the TMS intervention. These assessments were conducted by the same experienced neurologist.

MDS-UPDRS-III scores were further divided into several subscores according to different items ^48,49^. Axial scores were the sum of items 3.1 and 3.9-3.13. Tremor scores were the sum of items 20a-e and 21a-b. Rigidity scores were the sum of items 22a-e. Bradykinesia scores were the sum of items 23a-b, 24a-b, 25a-b, and 26a-b. Gait scores were directly from item 29.

### MRI preprocessing

The processing of both rs-fMRI and structural data was conducted using the pBFS Cloud v1.0.7 (Neural Galaxy Inc., Beijing). The preprocessing pipeline, primarily developed from our previously described pipeline ^25,50,51^, was adapted with software substitutions. The fMRI preprocessing sequence encompassed the following steps: (1) slice timing correction through stc_sess from the FreeSurfer version 6.0.0 software package (http://surfer.nmr.mgh.harvard.edu), (2) head motion correction using mc_sess from FreeSurfer (https://surfer.nmr.mgh.harvard.edu/fswiki/mc-sess), (3) linear detrending and bandpass filtering within the range of 0.01-0.08 Hz, and (4) regression to account for nuisance variables, which encompassed the six motion parameters, white matter signal, ventricular signal, global signal, and their first-order temporal derivatives.

Given the background noise in MP2RAGE T1w images in the PD dataset, the brain was first extracted from the uniform T1-weighted image using the ANTs. The subsequent preprocessing steps are consistent across structural sequences from the three datasets. The FreeSurfer version 6.0.0 software package (http://surfer.nmr.mgh.harvard.edu) was employed for processing ^52^. Surface mesh representations of the cerebral cortex were reconstructed from T1w images and non-linearly aligned to a shared spherical coordinate system. The functional and structural images were coregistered using boundary-based affine registration from the FsFast software package (http://surfer.nmr.mgh.harvard.edu/fswiki/FsFast). For the surface preprocessing pipeline, the functional images were aligned with the FreeSurfer cortical surface template (fsaverage6, 40,962 vertices per hemisphere). Applying a 6-mm full-width half-maximum (FWHM) surface smoothing kernel, the fMRI data was smoothed on the cortical surface. For the volumetric preprocessing pipeline, the preprocessed functional images in the native space were normalized to a 2-mm spatial resolution volumetric template (the FSL-version MNI ICBM152 nonlinear template) using coregistration matrix and the volumetric nonlinear registration facilitated by the Advanced Normalization Tools (ANTs) ^53^. Subsequent to the normalization step, a 6-mm FWHM isotropic smoothing Gaussian kernel was applied to the registered fMRI data within the brain mask.

### Seed-based RSFC analyses

In this study, we conducted three kinds of seed-based RSFC analyses. First, we performed whole-brain RSFC analyses employing seeds derived from both the SCAN and effector-specific networks. Second, we executed cortical RSFC analyses using the sweet spots of conventional DBS targets as seeds. Last, we investigated the RSFC between PD stimulation targets and the SCAN. To estimate the seed-based RSFC maps, we calculated Pearson correlation between the average BOLD fMRI signals within the seed ROI and the signals from cortical vertices or whole-brain voxels for each participant. Subsequently, we converted the correlation coefficients (r values) into z values through Fisher’s r-to-z transformation, normalizing the correlation coefficients. To generate group-averaged RSFC maps, we calculated the mean of the individualized z-maps across all participants within each patient or healthy participant group. To compare the SCAN-subcortical RSFC strength between PD patients and healthy participants, we used the 90^th^ percentile in RSFC strength within each subcortical structure, such as the striatum and cerebellum, when calculating SCAN-subcortical RSFC strength. ROIs for the STN-DBS targets were generated using the VTA of each patient, as detailed in the subsequent subsection. The TMS target was represented by a 6- mm spherical ROI, centered at the target coordinates defined in MNI space (-6, -6, 77) ^3^. Beyond the visualization of the seed-based RSFC maps, we computed the average RSFC across voxels within subregions and sweet spot ROIs (Fig. 3) or vertices within the SCAN or other effector-specific networks (Fig. 4) to quantify differences in RSFC among various networks.

### Identification of the SCAN

To identify the SCAN, we performed a two-stage analysis that includes an exploration stage and a network identification stage. First, to explore the existence of both individualized and group-level SCAN, we placed a continuous line of seeds along the precentral gyrus and estimated their RSFC. Second, the identification of the SCAN, along with other effector-specific networks such as those related to the foot, hand, and mouth, was accomplished using a fine-grained parcellation comprising a total of 213 cortical regions in both cortical hemispheres. This parcellation scheme, as previously detailed ^25,41,54^, was designed following the ‘divide and conquer’ principle. Each hemisphere was divided into five distinct lobes or regions, including the frontal, parietal, temporal, occipital lobes, and central region (precentral and postcentral gyri), according to the Desikan- Killiany atlas ^55^. Employing a k-means clustering algorithm, each lobe was parcellated into multiple cortical regions based on group-averaged RSFC profiles across 1,000 participants from the Genomic Superstruct Project (GSP) ^56^. The RSFC profile was determined as the connectivity between the BOLD signals of individual vertices from the FreeSurfer fsaverage6 surface and 1,175 ROIs uniformly distributed throughout the cerebral cortex ^57^. Identification of the SCAN and the three effector-specific networks within the fine-grained parcellation was based on a combination of anatomical landmarks and RSFC patterns. To further establish individual-specific fine-grained parcellations, an iterative parcellation strategy previously reported was applied ^58^. Through this iterative process, individualized parcellations for each lobe were derived, delineating the individual-specific SCAN and effector-specific networks for each participant.

### DBS electrode localization and the VTA estimation

Presurgical T1w MRI scans and postsurgical CT images were used to localize the electrodes. This procedure, akin to that outlined by Horn & Kühn ^59^, involved co-registering postsurgical CT images with presurgical T1-weighted images through linear registration using SPM12 (Wellcome Department of Cognitive Neurology, London, UK). Both CT and presurgical T1-weighted images were subsequently normalized to the MNI ICBM152 template using advanced normalization tools (ANTs). Semi-automated identification of DBS electrode contacts was then carried out on normalized CT images. The reconstruction of DBS electrodes from all 14 patients and various subcortical nuclei was achieved in MNI space using the LEAD-DBS software^45^.

Estimation of the VTA followed a previously established procedure ^60^. This process entailed generating a tetrahedral volume mesh based on the surface mesh of DBS contacts and subcortical nuclei using the Iso2Mesh toolbox within the LEAD-DBS software. Different regions were modeled as containing electrode materials, gray matter, or white matter. Conductivity values of 0.33 S/m and 0.14 S/m were assigned to gray and white matter, respectively. For platinum/iridium contacts and insulated electrode segments, values of 108 S/m and 10216 S/m were employed, respectively. Using the volume conductor model, the potential distribution stemming from DBS was simulated through the integration of the FieldTrip-SimBio pipeline. The applied voltage to active electrode contacts served as a boundary condition. Subsequently, the gradient of the potential distribution was computed through finite element method (FEM) derivation. The resulting gradient, being piecewise continuous due to the application of first-order FEM, was then thresholded for magnitudes surpassing the commonly used threshold of 0.2 V/mm. This delineated the extent and configuration of the VTA.

### TMS-induced E-field modeling

The TMS induced electric field (E-field) simulation was performed for each patient from the active group of the TMS dataset using SimNIBS ^61^ and the MagStim D70 coil model, following previous reports ^25,62^. The stimulation sites included both the left SMA target as well as the commonly-used TMS targets, i.e., hand- and foot-specific M1 ^22^. Specifically, the coil was placed tangentially to the scalp, with its center point directly over the stimulation site. Its orientation was posterior, following the direction towards the F2 landmark for the SMA target and FCz for the M1 targets in the 10-20 EEG system. For all stimulations, we opted for the fixed value of dI/dt = 1 A/µs, given it does not impact the spatial distribution of the E-field. Next, we generated a thresholded E-field map as the 99^th^ and 95^th^ percentile strongest E-field ^25,62,63^. Finally, we overlapped the individual thresholded E-field maps from each group to obtain an E-field overlap map.

### Statistical analysis

Statistical analyses were conducted utilizing the Scipy (v1.7.3) statistical package in Python. To assess group differences across various domains, two-tailed two-sample t-tests were employed for age, education, clinical symptom scores, imaging quality measures (relative head motion and tSNR), and RSFC. Chi-squared tests were used to examine differences in gender distribution among groups. Furthermore, two-tailed paired-sample t-tests were conducted to evaluate differences in RSFC between different functional networks as well as changes in RSFC and clinical scores before and after the intervention in PD patients. To investigate potential relationships between clinical scores and functional connectivity while accounting for covariates such as age and sex, partial Pearson correlations were performed. To address the issue of multiple comparisons, we applied the false discovery rate (FDR) correction method.

## Acknowledgement

We thank Yi Li for drawing illustrative figures (Fig. 1e, Fig. 5a, and Extended Data Fig. 6) and Ye Tian and Feng Zhang for the identifying electrodes’ location. This work was supported by the Changping Laboratory (H.L.), the China Postdoctoral Science Foundation (2022M720529 to J.R.), the National Natural Science Foundation of China (81527901 to L.L., 81720108021 to M.W., 81971689 to G.J., 31970979 to K.W., and 82090034 to K.W.), the National Key R&D Program of China (2017YFE0103600 to M.W.), and the Collaborative Innovation Center of Neuropsychiatric Disorders and Mental Health of Anhui Province (2020xkjT05 to K.W.).

## Competing interests

H.L. is the chief scientist of Neural Galaxy Inc. L.L. serves on the scientific advisory board for Beijing Pins Medical Co., Ltd and are listed as inventors in issued patents and patent applications on the deep brain stimulator used in this work. Other authors declare no conflict of interest regarding the publication of this work.

---

**Extended Data Table 1.** Characteristics of the PD, DBS, and TMS datasets.

| Datasets |  | PD dataset |  |  | p-val | DBS dataset |  | p-val | TMS dataset |  | p-val |
| --- | --- | --- | --- | --- | --- | --- | --- | --- | --- | --- | --- |
|  |  | Patient all | Patient | Control |  | Patient | Control |  | Active | Sham |  |
| demographics | Sample size | 166 | 65 | 60 | - | 14 | 25 | - | 19 | 19 | - |
|  | Sex | 64F/102M | 28F/37M | 34F/26M | 0.18<sup>a</sup> | 5F/9M | 13F/12M | 0.52<sup>a</sup> | 8F/11M | 6F/13M | 0.74<sup>a</sup> |
|  | Age<br>mean(std) | 61.81(7.84) | 58.57(8.57) | 56.10(6.64) | 0.08<sup>b</sup> | 54.71(7.65) | 56.32(6.88) | 0.50<sup>b</sup> | 61.37(7.33) | 59.74(8.86) | 0.27<sup>b</sup> |
|  | Education (yr)<br>mean(std) | 8.60(3.75) | 8.33(3.84) | 9.92(3.68) <sup>c</sup> | 0.03<sup>b,c</sup> | 10.90(3.00) | - | - | 8.64(5.36) | 8.42(4.51) | 0.44<sup>b</sup> |
|  | Disease duration (yr)<br>mean(std) | 5.82(4.08) | 5.11(2.80) | - | - | - | - | - | 4.16(2.44) | 5.29(3.81) | 0.86<sup>b</sup> |
| clinical information | UPDRSIII | 38.37(19.86) | 38.95(19.45) | - | - | 32.41(13.13) | - | - | 27.05(10.15) | 29.68(9.14) | 0.80<sup>b</sup> |
|  | H-Y | 2.35(0.91) | 2.31(1.02) | - | - | 3.5(0.8) | - | - | 1.63(0.62) | 1.71(0.48) | 0.67<sup>b</sup> |
|  | MMSE | 25.57(4.15) | 25.65(3.78) | 27.47(2.53) | 0.002<sup>b</sup> | 27.7(1.4) | - | - | 27.89(2.02) | 28.63(1.86) | 0.87<sup>b</sup> |
|  | HAMD | 10.78(5.79) | 11.86(6.26) | 2.77(2.18) | <0.001<sup>b</sup> | - | - | - | 7.84(4.90) | 7.42(5.67) | 0.40<sup>b</sup> |
|  | HAMA | 10.73(6.22) | 12.49(6.93) | 2.3(2.02) | <0.001<sup>b</sup> | - | - | - | 8.53(5.47) | 7.95(5.60) | 0.37<sup>b</sup> |
| imaging quality | Head motion (mm)<br>mean(std) |  | 0.09(0.04) | 0.11(0.03) | 0.99<sup>b</sup> | 0.10(0.04) | 0.10(0.04) | 0.43<sup>b</sup> | 0.03(0.01) | 0.04(0.02) | 0.92<sup>b</sup> |
|  | SNR<br>mean(std) |  | 36.39(4.09) | 35.88(3.34) | 0.22<sup>b</sup> | 72.19(9.84) | 73.42(7.96) | 0.66<sup>b</sup> | 65.70(9.06) | 68.37(10.63) | 0.79<sup>b</sup> |

<sup>a</sup>: Chi-square test<sup>b</sup>: Independent t-test<sup>c</sup>: Mean and STD were calculated based on 50 healthy people.

**Extended Data Table 2.** Characteristics of DBS sweet spots, VIM-MRgFUS, and STN-segmentation datasets.

| Datasets | Sample size | Sex | Age (yr)<br>mean±std | Disease duration (yr)<br>mean±std | Disease severity<br>(clinical scale)<br>mean±std | Clinical improvement<br>at Follow-up<br>mean±std | references |
| --- | --- | --- | --- | --- | --- | --- | --- |
| Sweet spots of STN-DBS | 275 PD | 80F/195M | 59.8±7.1 | 11.5±4.4 | 36.5±11.1<br>(UPDRS-III) | 51.2±24.8% | Elias et al.<sup>14</sup> |
| Sweet spots of GPi-DBS | 28 PD | 13F/15M | 64.4±7.0 | 14.8±5.7 | 39.9±14.0<br>(UPDRS-III) | 38.9±27.0% | Elias et al.<sup>14</sup> |
| Sweet spots of VIM-DBS | 33 ET/6 others | 13F/26M | 64.3±11.6 | 28.8±18.1 | 58.7±11.3<br>(TRS) | 39.7±22.6% | Elias et al.<sup>14</sup> |
| Lesion overlap of VIM-MRgFUS | 10 PD | 2F/8M | 55.4±7.2 | 5.2±1.7 | 14.6±5.9<br>(CRST Part A) | 41.4±18.1% | Dahmani et al.<sup>2</sup> |
| STN segmentation | 13 HC | 6F/7M | 50.6±10.9 | - | - | - | Accolla et al.<sup>15</sup> |

**Extended Data Fig. 1.**
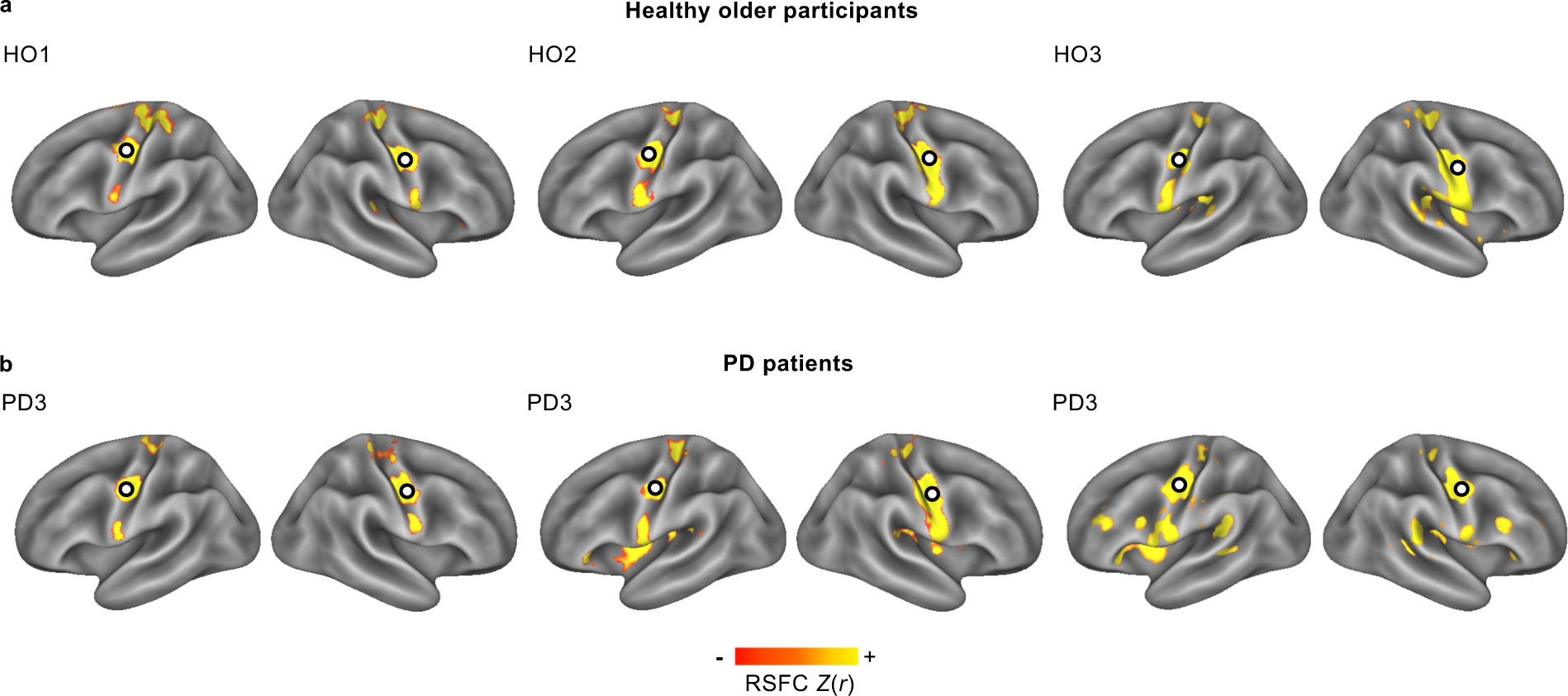
The SCAN is detectable in three additional healthy older individuals and PD patients. The characteristic SCAN motif is observed in (a) three healthy older participants (HO1-HO3, age > 65 years) and (b) three PD patients (PD1-PD3). Circles indicate the seed regions of interest located in the middle regions of the SCAN.

**Extended Data Fig. 2.**
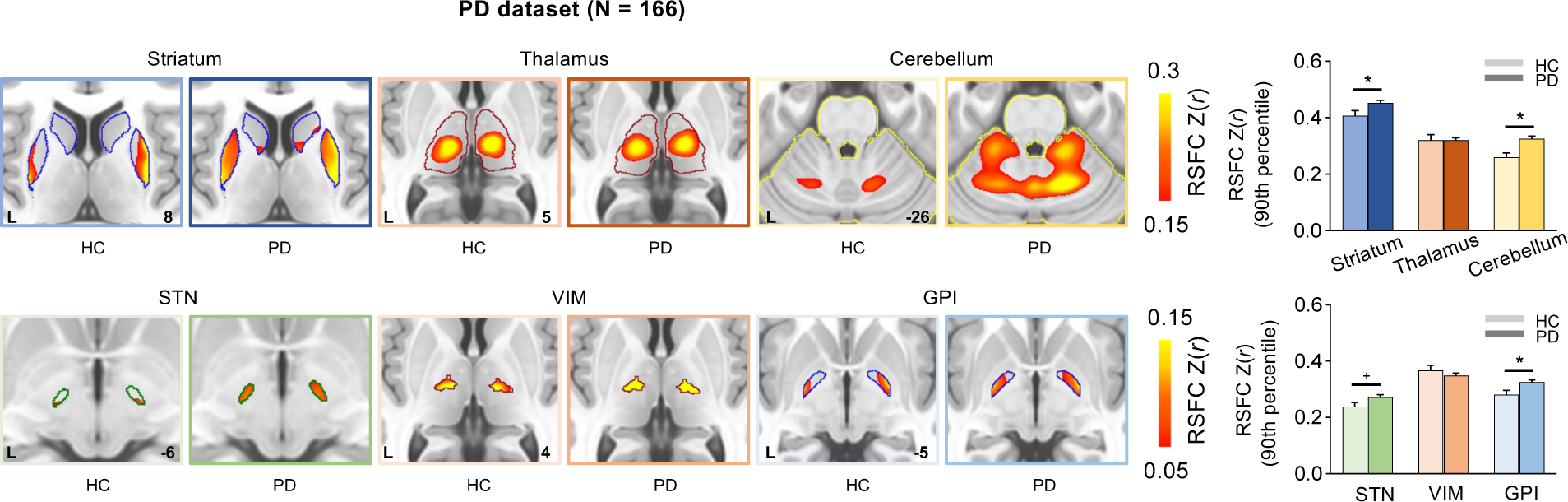
Replication of abnormal hyper-connectivity between SCAN and subcortical regions in full PD dataset. The analysis shown in the Fig. 1 is replicated in 166 PD patients from the PD dataset, with PD patients demonstrating statistically stronger RSFC in the striatum, cerebellum, and GPi (*p values < 0.01, FDR-corrected), trend towards greater RSFC in the STN (+p = 0.052, FDR-corrected) than healthy controls.

**Extended Data Fig 3.**
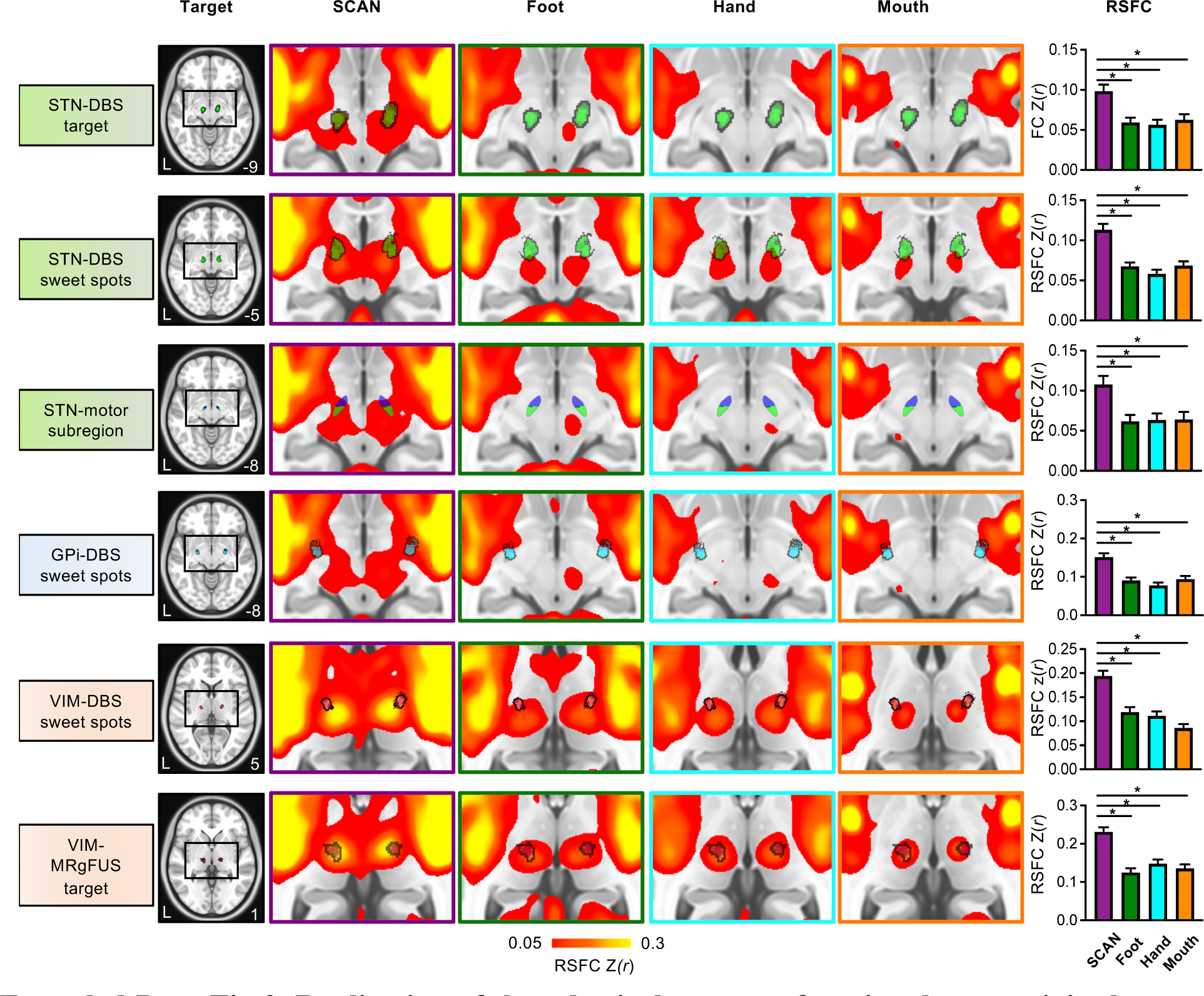
Replication of the selectively greater functional connectivity between SCAN and diverse neuromodulatory targets in the full PD dataset. The analysis presented in the Fig. 3 is replicated in a cohort of 166 PD patients from the PD dataset, revealing selectively greater functional connectivity between SCAN and diverse neuromodulatory targets (all p values < 0.01, FDR-corrected).

**Extended Data Fig. 4.**
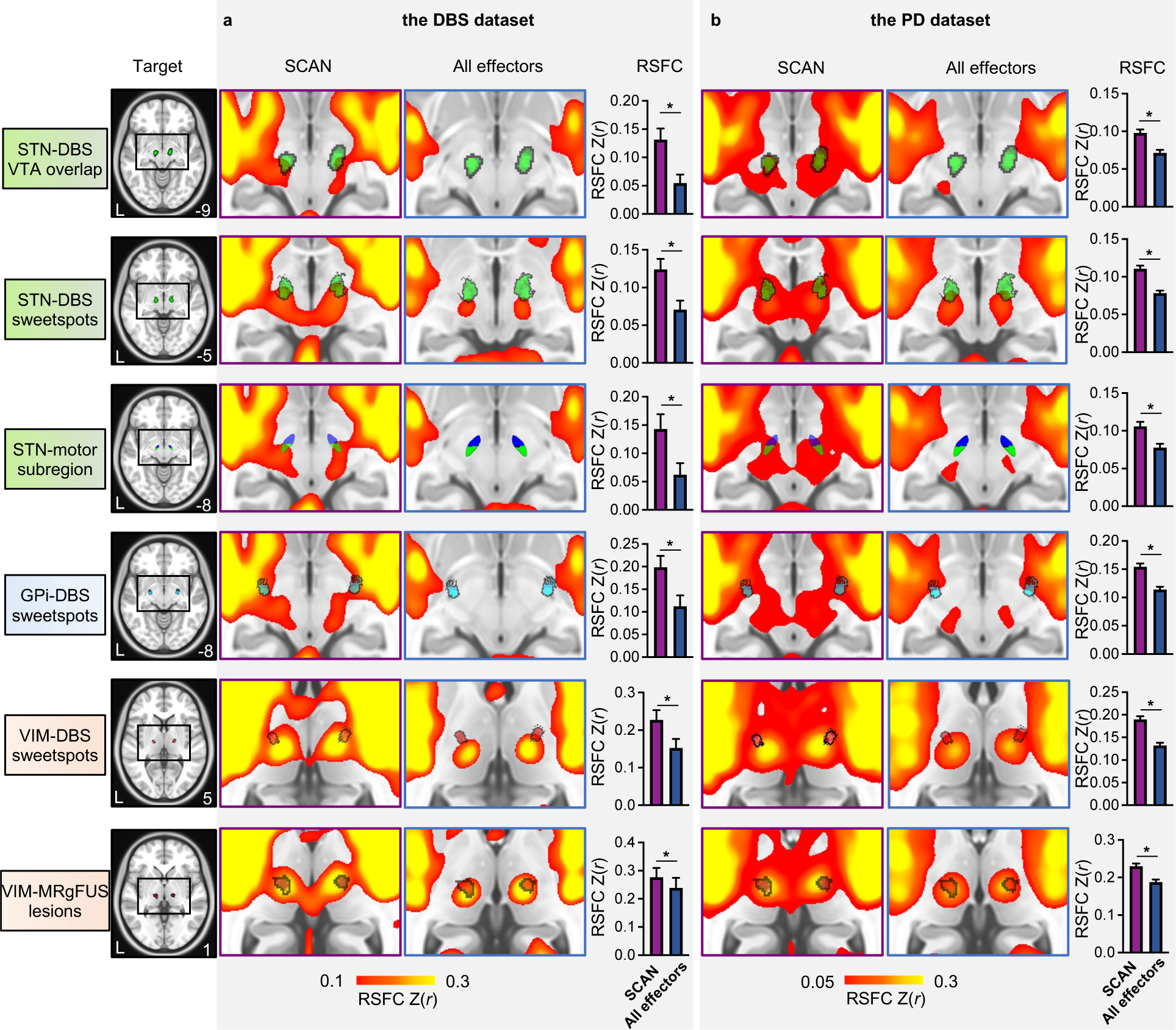
The SCAN shows stronger RSFC with neuromodulatory targets than the combinations of effector-specific regions. To address potential confounding influences associated with the composite nature of the SCAN, which encompasses three distinct regions, as opposed to the single region within each effector-specific network, we performed a comparative analysis of RSFC targeting the SCAN versus RSFC targeting an average of the three effector-specific networks. Across both the (a) DBS and (b) PD datasets, the SCAN-target RSFC displayed significant enhancement (all p-values < 0.01, FDR-corrected).

**Extended Data Fig. 5.**
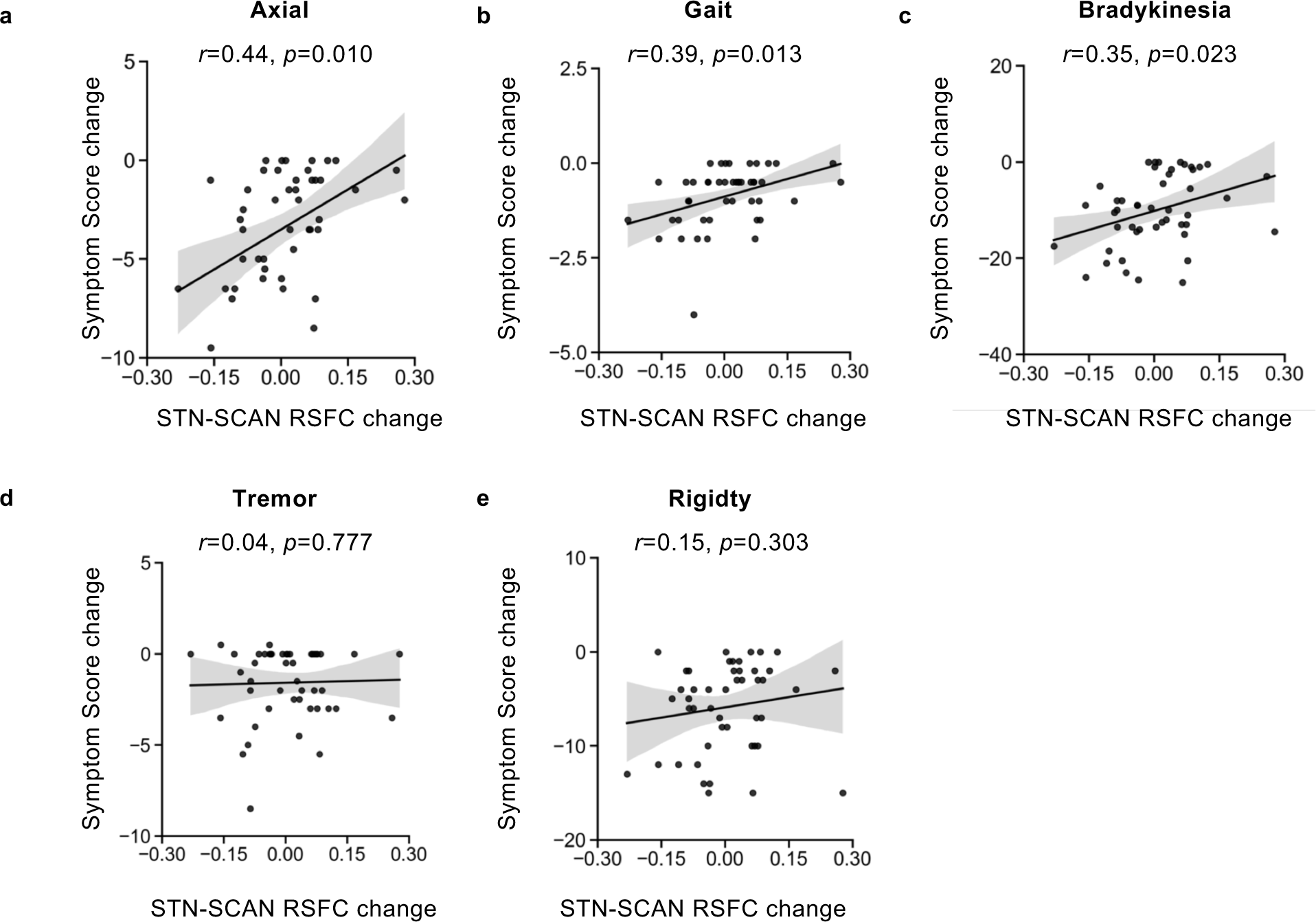
Changes in target-SCAN RSFC induced by STN-DBS are associated with changes in various motor symptoms. The RSFC changes show significant associations with (a) axial movement (r = 0.44, p = 0.010, FDR-corrected), (b) gait (r = 0.39, p = 0.013, FDR-corrected), and (c) bradykinesia (r = 0.35, p = 0.023, FDR-corrected), but showed negligible correlations with (d) tremor (r = 0.04, p = 0.777, FDR-corrected) and (e) rigidity (r = 0.15, p = 0.303, FDR-corrected).

**Extended Data Fig. 6.**
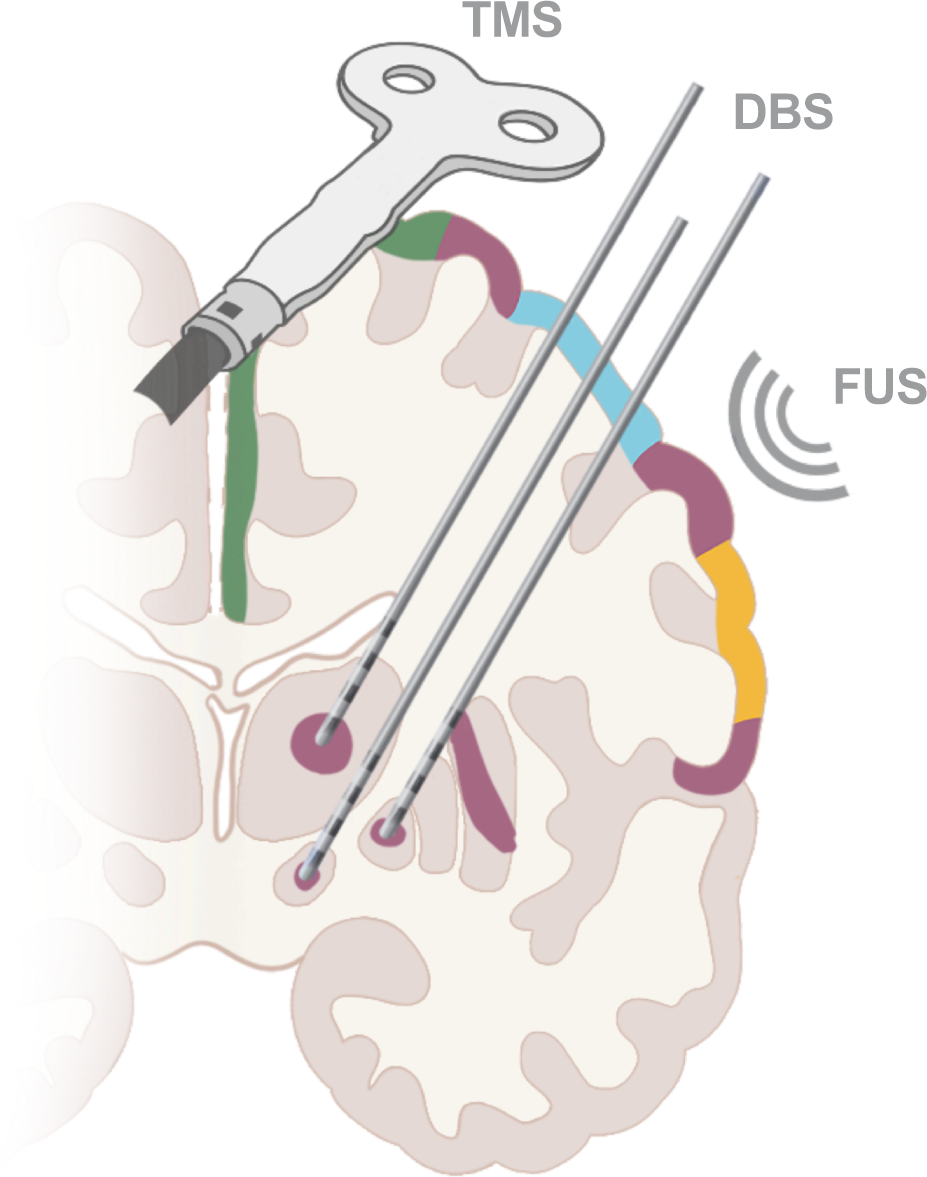
The illustration depicts multiple types of neuromodulation targeting the SCAN for PD treatment. The cortical and subcortical regions in the SCAN are represented in purple. Various neuromodulation techniques, including TMS, DBS, and FUS, targeting the SCAN, hold the potential for alleviating PD symptoms.

## Supplementary Information

**Supplementary Fig. 1.**
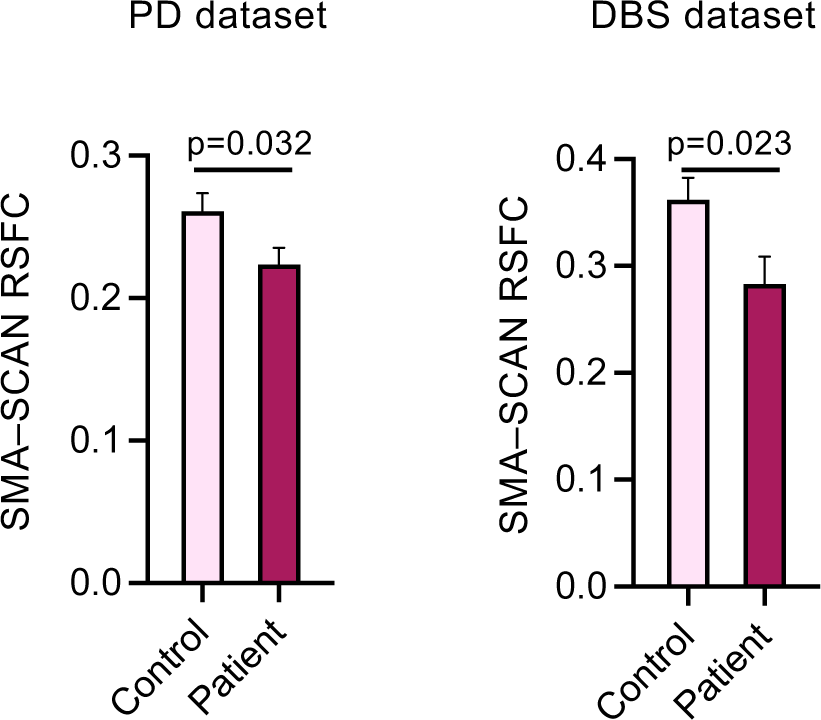
Reduced SMA-SCAN RSFC strength in PD patients compared to healthy participants. The RSFC between the left supplementary motor area (SMA) target of rTMS (MNI coordinates: -6, 6, 77) and the SCAN was estimated in PD patients and healthy participants in both PD and DBS datasets. SMA-SCAN RSFC is significantly weaker in PD patients when compared to aged healthy participants (two-sample paired t-tests, t(123) =2.17 , p = 0.032 in the PD dataset; t(37) = 2.37, p =0.023 in the DBS dataset).

**Supplementary Fig. 2.**
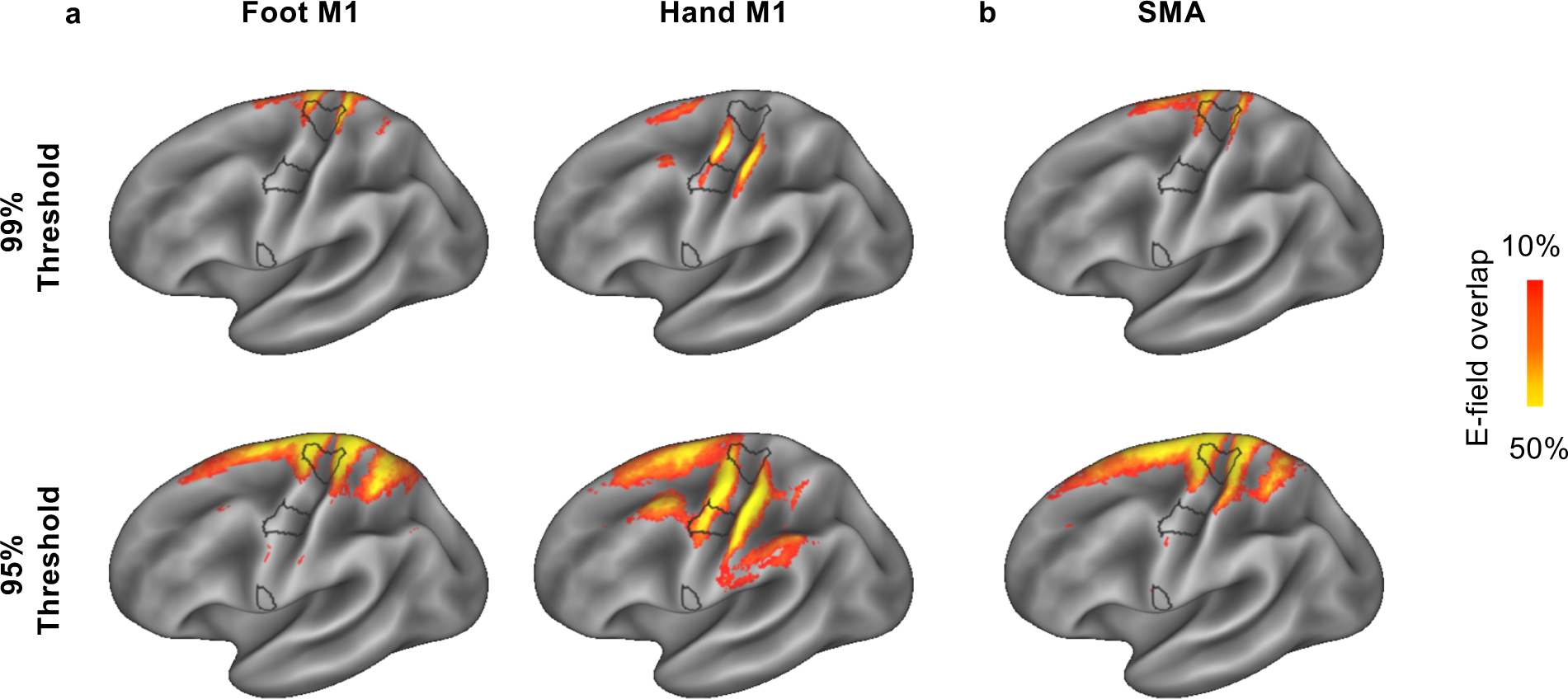
Partial overlapping E-fields of conventional rTMS targets for PD partially overlaps with the SCAN. Conventional rTMS targets for PD include hand/foot M1 and SMA. E-field map was estimated on each subject’s individual surfaces and projected onto a common surface to evaluate the overlap among subjects. The thresholded E-field maps, representing the 99th (top panel) and 95th (bottom panel) percentile of the strongest E-field values, reveal partial overlap with the SCAN (outlined in black) in (a) the Hand/Foot M1 targets and (b) the SMA target.

**Supplementary Table 1.**
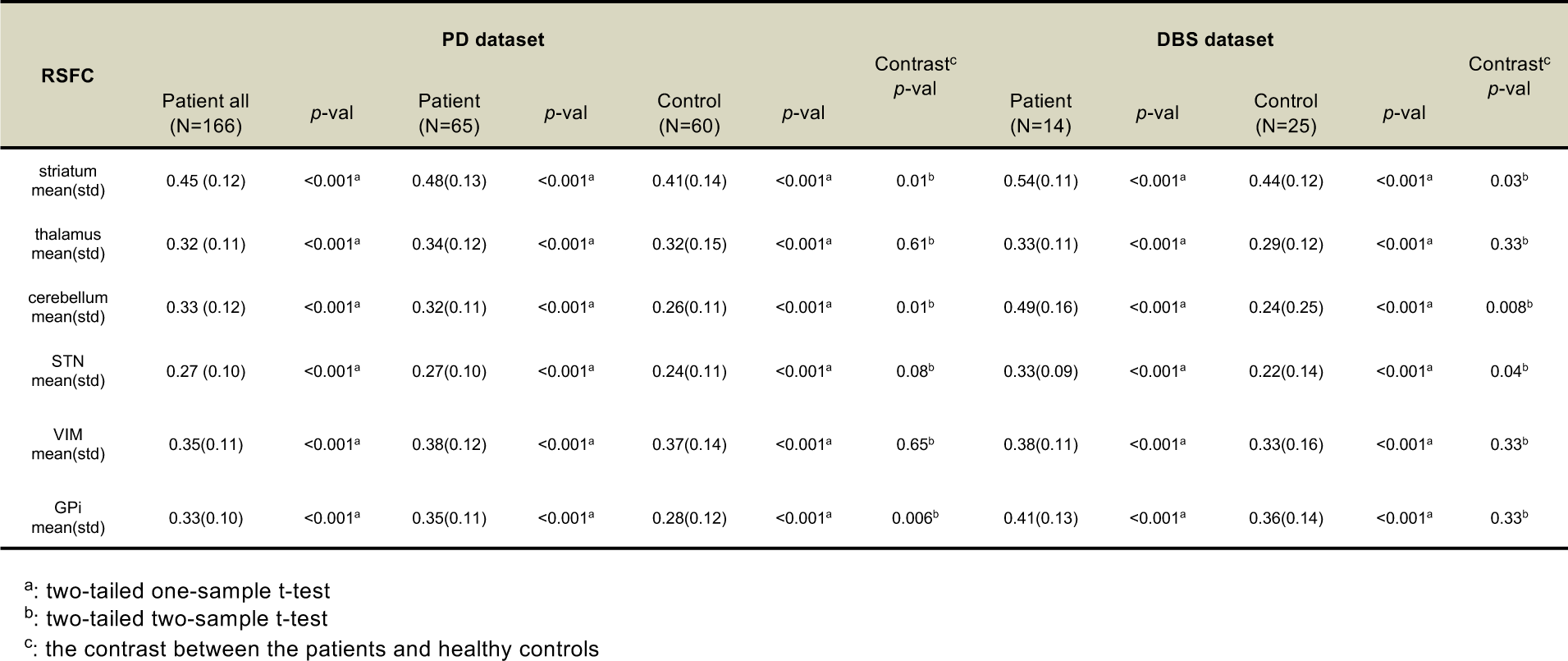
All RSFC strengths and statistical results.

**Supplementary Table 2.** STN-DBS Stimulation Parameters.

| Patient ID | 1-month |  | 3-month |  | 6-month |  | 12-month |  |
| --- | --- | --- | --- | --- | --- | --- | --- | --- |
|  | Amplitude (V) / Pulse width (μs) |  | Amplitude (V) / Pulse width (μs) |  | Amplitude (V) / Pulse width (μs) |  | Amplitude (V) / Pulse width (μs) |  |
|  | Left | Right | Left | Right | Left | Right | Left | Right |
| DBS01 | 1.5/60 | 1.7/60 | NA | NA | NA | NA | NA | NA |
| DBS02 | 2.1/60 | 2.1/60 | 2.5/60 | 2.5/70 | 2.6/60 | 2.5/60 | 2.6/60 | 2.5/60 |
| DBS03 | 1.5/60 | 1.8/60 | 2.1/60 | 2.0/60 | NA | NA | NA | NA |
| DBS04 | 1.5/60 | 2.0/60 | 2.2/60 | 2.2/60 | 1.7/60 | 3.1/60 | 1.7/60 | 3.1/60 |
| DBS05 | 1.9/60 | 1.8/80 | 2.3/60 | 2.3/60 | 2.7/90 | 3.0/60 | 2.7/90 | 3.0/60 |
| DBS06 | 2.5/80 | 1.9/60 | 2.5/70 | 2.5/60 | 2.8/70 | 2.9/60 | 2.8/70 | 2.9/60 |
| DBS07 | 1.9/60 | 1.4/60 | 2.6/80 | 2.4/70 | 2.9/70 | 2.7/60 | 2.9/70 | 2.7/60 |
| DBS08 | 1.7/60 | 1.8/60 | 1.6/60 | 1.7/60 | 1.3/50 | 1.2/60 | 1.3/50 | 1.2/60 |
| DBS09 | 1.9/60 | 2.5/60 | 1.9/60 | 2.5/60 | 2.5/60 | 2.7/60 | 2.5/60 | 2.7/60 |
| DBS10 | 1.0/60 | 1.9/60 | 1.0/60 | 2.4/60 | 1.1/60 | 2.1/60 | 1.1/60 | 2.1/60 |
| DBS11 | 2.0/80 | 1.8/60 | 2.0/70 | 1.8/70 | 3.6/90 | 3.5/80 | 3.7/90 | 3.8/70 |
| DBS12 | 2.0/60 | 2.0/60 | 2.2/60 | 2.4/60 | 2.5/60 | 2.4/60 | 2.8/50 | 2.6/60 |
| DBS13 | 2.5/80 | 2.2/40 | 2.5/80 | 2.5/60 | 3.0/70 | 2.7/60 | 3.2/80 | 3.1/60 |
| DBS14 | 1.3/60 | 1.5/60 | 1.7/60 | 2.6/60 | 2.1/60 | 2.2/70 | 2.4/60 | 2.8/60 |
| Mean | 1.8/64.3 | 1.9/60.0 | 2.1/64.6 | 2.3/62.3 | 2.4/66.7 | 2.6/62.5 | 2.5/66.7 | 2.7/60.8 |
| Std | 0.4/8.5 | 0.3/7.8 | 0.5/7.8 | 0.3/4.4 | 0.7/12.5 | 0.6/6.2 | 0.8/13.7 | 0.6/2.9 |

## Notes

### Competing Interest Statement

H.L. is the chief scientist of Neural Galaxy Inc. Luming Li serve on the scientific advisory board for Beijing Pins Medical Co., Ltd and are listed as inventors in issued patents and patent applications on the deep brain stimulator used in this work. Other authors declare no conflict of interest regarding the publication of this work.

